# Structural basis for surface activation of the classical complement cascade by the short pentraxin C-reactive protein

**DOI:** 10.1101/2024.03.18.585147

**Authors:** Dylan P. Noone, Marjolein M. E. Isendoorn, Sebastiaan M. W. R. Hamers, Mariska E. Keizer, Jip Wulffelé, Tijn T. van der Velden, Douwe J. Dijkstra, Leendert A. Trouw, Dmitri V. Filippov, Thomas H. Sharp

## Abstract

Human C-reactive protein (CRP) is a pentameric complex involved in defence against pathogens and regulation of autoimmunity. CRP is also a therapeutic target, with both administration and depletion of serum CRP being pursued as a possible treatment for autoimmune and cardiovascular diseases, among others. CRP binds to phosphocholine (PC) moieties on membranes in order to activate the complement system via the C1 complex, but it is unknown how CRP, or any pentraxin, binds to C1. Here, we present a cryo-electron tomography (cryoET)-derived structure of CRP bound to PC ligands and the C1 complex. To gain control of CRP binding, a synthetic mimotope of PC was synthesised and used to decorate cell-mimetic liposome surfaces. Structure-guided mutagenesis of CRP yielded a fully-active complex able to bind PC-coated liposomes that was ideal for cryoET and subtomogram averaging. In contrast to antibodies, which form Fc-mediated hexameric platforms to bind and activate the C1 complex, CRP formed rectangular platforms assembled from four laterally-associated CRP pentamers that bind only four of the six available globular C1 head groups. Potential residues mediating lateral association of CRP were identified from interactions between unit cells in existing crystal structures, which rationalised previously unexplained mutagenesis data regarding CRP-mediated complement activation. The structure also enabled interpretation of existing biochemical data regarding interactions mediating C1 binding, and identified additional residues for further mutagenesis studies. These structural data therefore provide a possible mechanism for regulation of complement by CRP, which limits complement progression and has consequences for how the innate immune system influences autoimmunity.

**Significance statement:** Human C-reactive protein (CRP) activates the complement system to protect us from infections, but can also contribute towards progression of cardiovascular and autoimmune diseases when erroneously activated. To understand these processes, the authors used cryo-electron tomography to solve the *in situ* structure of surface-bound CRP interacting with the complement C1 complex. The structure revealed new interfaces that explain previous, sometimes contradictory, biochemical data. Comparisons with existing structures of antibody-mediated C1 activation revealed distinct structural differences that may explain how CRP modulates complement activity. Together, these structural data identify residues for mutagenesis to gain control over CRP functions, and provide new routes for future therapeutic developments.

## Introduction

C-reactive protein (CRP) is an acute phase protein of the pentraxin family, and is produced by the liver in response to inflammatory stimuli such as interleukin-6 (1). CRP is found at sites of inflammation (2, 3), can activate proinflammatory pathways such as the classical complement pathway (4) and can agglutinate cells and cellular debris (5), which facilitates their phagocytosis (6, 7). Additionally, CRP is linked to the pathogenesis of cardiovascular disorders such as atherosclerosis (8), with CRP serum levels directly correlating with patient prognosis (9). CRP also helps regulate autoimmunity, being linked to autoimmune disorders such as systemic lupus erythematosus (SLE) (10), and administration of CRP to mouse models of SLE results in the reversal of symptoms (11, 12), suggesting that CRP has a protective role.

CRP circulates in human serum as a disc-like pentameric complex of 115 kDa, consisting of non-covalently associated monomers that are linked via a network of ionic bonds (13-15). The pentamer has two faces; the binding face (B-face) and the activating face (A-face) (16). The B-face of each CRP monomer coordinates two Ca^2+^ ions, which are important for binding to the phosphocholine (PC) headgroup of phosphatidylcholine lipids displayed on membranes(5, 17). Phosphatidylcholine is found in both human cells (18) and pathogens (1), reflecting the role of CRP in both the clearance of apoptotic cells (19) and innate immune defence (20). CRP can freely associate with synthetic PC-containing conjugates (21), but can only bind to lipid membranes under the conditions of lipid oxidation (21), high membrane curvature (22) or in the presence of lysophosphatidylcholine (LPC). LPC is posited to disrupt the normal packing of lipid head groups in lipid bilayers (23) exposing the PC headgroup to enable CRP binding (24) as well as being associated with apoptosis (25) and disease states such as atherosclerosis (26).

The A-face of CRP associates with effector ligands such as Fcγ receptors (6, 7), which mediate phagocytosis (5). Pentameric CRP can also oligomerise into decamers via A-face stacking (15), which may cause agglutination of cells or cellular debris (5, 27). The initiating component of the classical complement pathway, the C1 complex (**Fig. S1**), also binds to the A-face before activation (28). The C1 complex contains C1q, a disulfide-linked hexamer of heterotrimeric proteins that form an N-terminal stalk which splays outwards to form 6 collagenous arms. The collagenous arms terminate in globular C1q (gC1q) domains that detect ligands, such as antibodies or pentraxins (29, 30). A heterotetrameric C1r_2_s_2_ protease complex is located between the C1q collagenous arms (31). Upon binding of gC1q to ligands, the proenzymes C1r and C1s are activated in turn, allowing C1s to cleave complement component C4 to form C4b. Formation of C4b exposes a reactive thioester bond that forms a covalent bond with adjacent molecules and surfaces. Next, C2 binds to C4b and is also cleaved by C1s, forming the C3-convertase C4b2b, which propagates the complement pathway (32).

IgG subclasses IgG1 and IgG3 are potent complement activators that circulate as monomers but form higher-order multimers on surfaces that are necessary to bind and activate the C1 complex (33, 34). In contrast, IgM circulates as a pentameric or hexameric complex, but must undergo a conformational change upon antigen binding to activate C1 (35). It is currently unclear whether similar mechanisms exist for CRP-mediated complement activation, and no structural data exists of CRP bound to membranes or activating complement. Furthermore, direct comparisons between CRP- and antibody-mediated complement activation show that antibodies result in greater lysis than CRP (36), and that CRP-mediated complement activation is more likely to result in opsonisation without membrane attack complex (MAC) pore formation, biasing the complement cascade away from lysis and towards phagocytosis and silent clearance (37). Understanding the activation mechanism by CRP may therefore reveal the physiological switch between phagocytosis or lysis, the former preventing the release of autoantigens from cellular debris, with consequences for the regulation of autoimmunity (38).

To investigate complement activation by CRP, we produced cell mimetic liposomes decorated with a synthetic version of PC via the linkage of PC-azide conjugates to lipids functionalised with dibenzocyclooctyne (DBCO) using click chemistry. CRP successfully activated the classical complement cascade on the cell mimetics. CRP also agglutinated the liposomes, forming large aggregates unsuitable for cryo electron tomography (cryoET) imaging and structural biology. We performed structure-guided rational modification of CRP to produce a single-point mutant of CRP that was incapable of agglutinating liposomes, but still able to activate complement. This mutant, with decoupled agglutination and complement activating abilities, could be used for cryoET of reconstituted PC-CRP-C1 complexes on liposomes. We observed lateral interactions between liposome-bound CRP, which formed a tetrameric complex that bound to and activated C1 and formed a structure distinct from antibody-mediated C1 binding. Additionally, the C1s_2_r_2_ protease platform adopted a pre-C4b cleavage conformation, with less well-defined density inhabiting a separate configuration, suggesting a possible mechanism for C1 autoactivation. These data provide structural details that explain how complement activation is biased between lysis or opsonophagocytosis by differential ligand binding.

## Results

### Structure-guided mutagenesis of CRP decouples agglutination and complement activation

To image CRP bound to C1 on a lipid membrane via cryoEM, we generated liposomes containing the native CRP ligand lysophosphatidylcholine (LPC). However, LPC alone appeared to be unstable, and liposomes containing LPC produced inconsistent results (**Fig. S2A**,**B**). Pure LPC formed ∼1 nm sized particles consistent with micelles and, whilst liposomes containing a portion of LPC appeared relatively homogenous in terms of size (**Fig. S2C**), they also presented as heterogenous populations of liposomes on cryoEM grids (**Fig. S2D**). We attributed this instability to the LPC, and so to gain more control over the system, and obtain homogenous and consistent liposome samples, a lipid formulation previously reported in the literature for the resolution of C1-antibody complexes (33-35) was used. To provide a ligand for CRP, we synthesised a PC compound with an azide moiety (**Fig. 1A**). This synthetic ligand could be attached to the liposomes by including a low concentration (5 mol%) of lipids displaying a dibenzocyclooctyne (DBCO)-modified headgroup, to which we covalently attached the azide-PC molecules using copper-free click chemistry. CRP bound to these liposomes and successfully activated the classical complement cascade comparably to liposomes containing LPC (**Figs. 1B & S3B**,**C**). However, successful imaging via cryoEM was hindered by the tendency of CRP to agglutinate the liposomes (**Fig. 1C**), which resulted in samples that were too thick for cryoET (39) and that also did not contain isolated liposomes decorated in protein bound to C1, as has been previously achieved for IgG1, IgG3 and IgM (33-35, 40).

**Figure 1:**
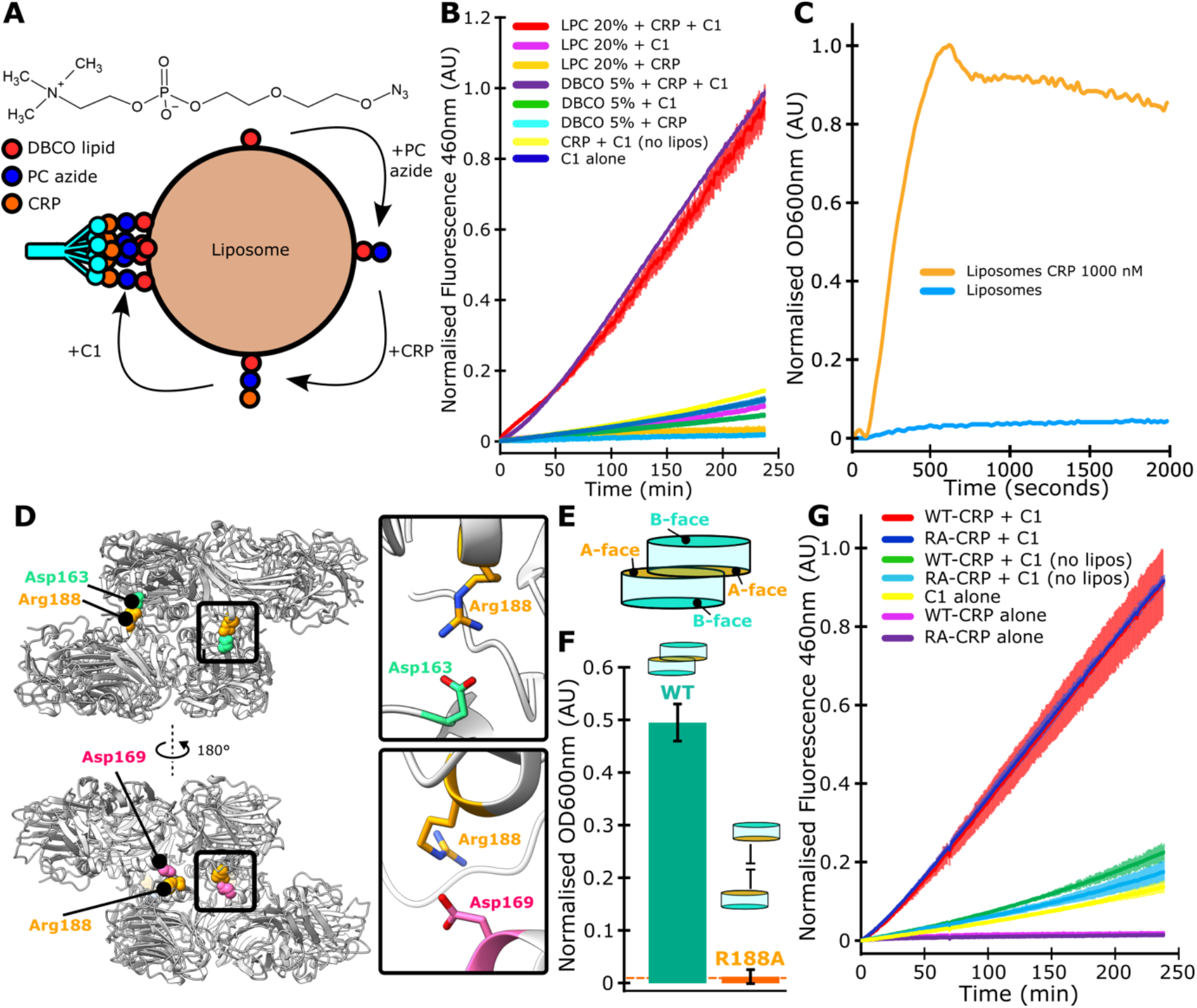
CRP mediated agglutination of liposomes is dependent on decamer formation. (**A**) Structure of PC-azide chemical (top), and a schematic of the strategy for in vitro reconstitution of PC-CRP-C1 complexes. (**B**) C1s activation assay of CRP bound to DBCO-functionalized liposomes conjugated to PC-azide, or LPC-containing liposomes. (**C**) Addition of CRP to liposomes caused an increase in the OD_600_. (**D**) CRP decamers (grey; PDB code 7PK9) contain four salt bridges involving the A-face residue Arg188. (**E**) Schematic of offset wild-type CRP decamers formed through A-face association of CRP pentamers. (**F**) Mutant RA-CRP (R188A) does not aggregate liposomes compared to wild type (WT) CRP. (**G**) C1s activity stimulated by liposomes bound by either WT-or RA-CRP.

Calcium dependent aggregation of lipids is a known function of CRP (41, 42), which we postulated to be mediated by a decameric species of CRP (15), whereby two CRP pentamers associate in an offset stack through A-face interactions (**Figs. 1D**,**E**) (5, 15, 16). By examining structures of decameric CRP in the protein database (PDB; 3PVO, 3PVN, 7TBA and 7PK9) (5, 15, 16), we identified residue Arg188 as the predominant mediator of decamerization, as it forms four ionic bonds per decamer with residues Asp163 or Asp169 on the opposing pentamer (**Fig. 1D**). Additionally, Arg188 does not overlap with the putative C1 binding site (**Fig. S4A**,**B**). It was therefore mutated to an alanine in an effort to prevent formation of the decameric species without affecting C1 binding or complement activation, before being expressed and purified using PC-conjugated agarose (43). The R188A mutant (RA-CRP) drastically reduced fluid phase aggregation of liposomes as measured via OD_600_ (**Fig. 1F**), behaved in an identical manner to wild type CRP (WT-CRP) on SDS-PAGE and activated complement to the same extent as WT CRP on liposomes (**Figs. 1G & S4C**).

#### *In situ* structure of a CRP-C1 complex

Our single point mutant of CRP, RA-CRP, did not agglutinate liposomes on grids prepared for cryoEM (**Fig. S5**), and it was possible to obtain thin samples suitable for cryoET. Incubation of liposomes coated with RA-CRP with purified C1 resulted in the imaging of density consistent with the C1 complex bound to the membranes (**Figs. 2A & S6**). RA-CRP formed a layer ∼4 nm thick, with C1 binding producing complexes that stood ∼30 nm from the membrane. PC-CRP-C1 complexes displayed density consistent with the C1r_2_s_2_ protease platform oriented parallel to the membrane (**Figs. 2A & S6**), as well as density flanking the central regions of the C1r_2_s_2_ protease platform that connected to the CRP layer (**Fig. S6**).

**Figure 2:**
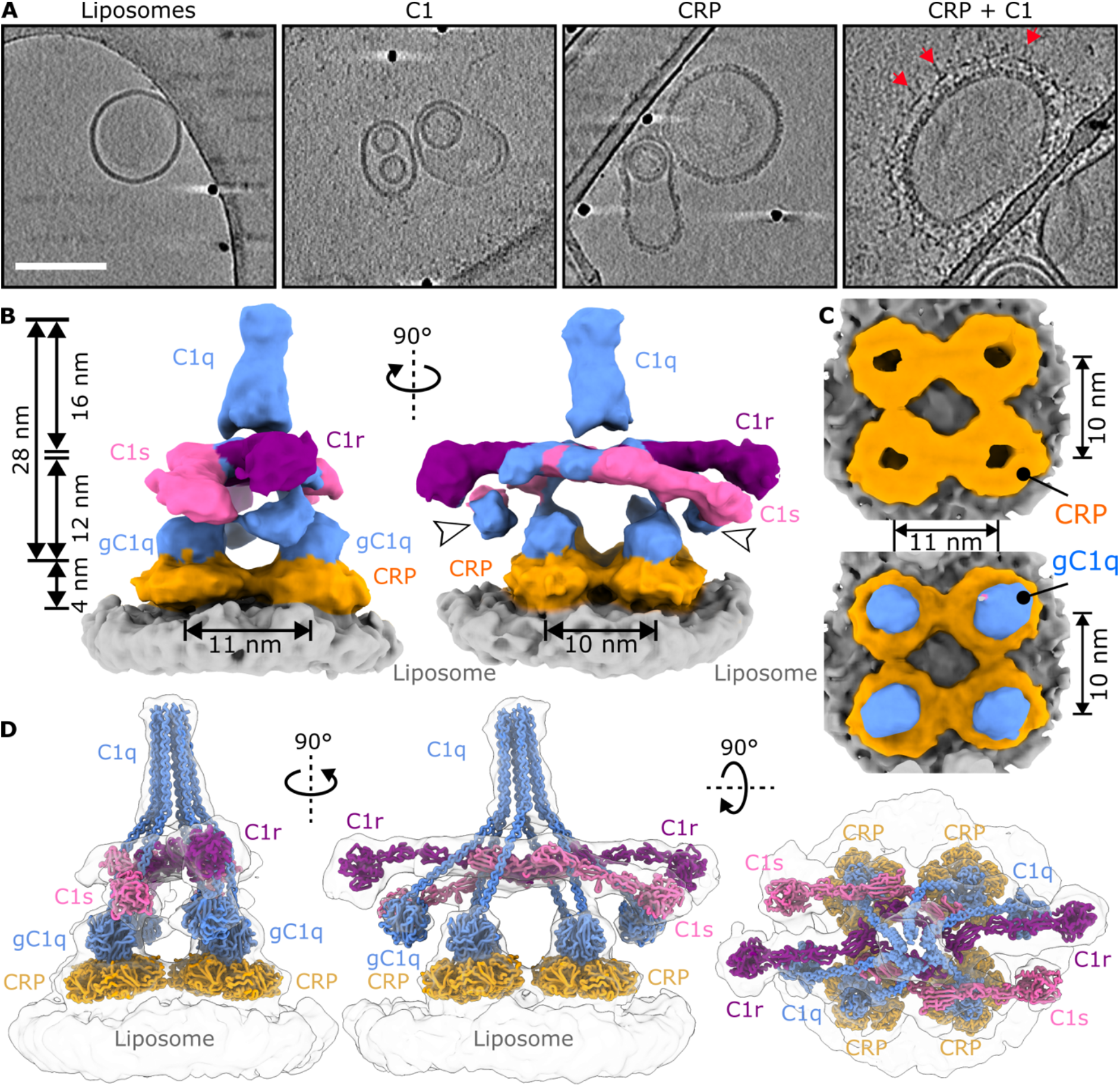
C1 binds to tetramers of CRP pentamers on lipid surfaces. (**A**) Slices 13.9 nm thick through cryotomograms showing CRP bound to liposomes, and the C1 complex on top (red arrows). Scale bar represents 100 nm. (**B**) Subtomogram map of the tetrameric CRP-C1 complex map lowpass filtered to 20 Å and coloured by protein. White arrowheads indicate gC1q domains not bound to CRP. (**C**) View through the tetrameric CRP platform (top) and with gC1q domains (bottom). (**D**) Fitted model with proteins annotated. Colouring: C1q, blue; C1r, purple; C1s, pink; CRP, orange; liposome, grey.

From 133 tomograms we were able to manually pick 3,428 particles of CRP-C1 complexes, which allowed us to perform subtomogram averaging (**Fig. S7**). After several rounds of refinement and 3D classification, 2,518 of the particles contributed to a map resolved to 24 Å (**Fig. S7**). The map contained identifiable CRP molecules, as well as all elements of the C1 complex (**Fig. 2B**). The map revealed four CRP pentamers with four associated gC1q domains bound (**Fig. 2C**). The map did not resolve the individual CRP monomers, and nor did multireference classification (**Fig. S8**). Instead, each CRP pentamer presented as a torus, indicating structural heterogeneity in this region. Two lobes of density were also present, ∼6.5 nm above the lipid bilayer, which corresponded to the remaining two gC1q domains that are not bound to a CRP pentamer (white arrowheads, **Figs. 2B**). Above these, the remainder of the C1 complex stood 28 nm tall, consistent with the measurements derived from the tomographic slices (**Fig. 2B**), with the C1r_2_s_2_ platform located between the C1q collagen arms, as seen in previously published models of C1 bound to antibodies (33-35).

### CRP pentamers interact laterally to form tetrameric platforms

Previously resolved C1-antibody complexes formed hexameric complexes (33-35, 40), yet CRP instead formed a tetrameric platform which bound to C1. Density corresponding to CRP protruded ∼4 nm from the membrane, with the centres of the tori arranged on a 10 × 11 nm rectangle (**Fig. 2 B,C**). The model built into the electron density (**Fig. 2D**), allowed closer inspection of the tetrameric CRP platform and interactions with the four gC1q domains.

CRP pentamers were placed in the toroidal densities with the known PC-binding B-face oriented towards the membrane. The low resolution of the map meant that determination of the rotational orientation of the pentamers was not possible. However, the map suggested lateral interactions between neighbouring CRP pentamers. To explore these lateral interfaces, we analysed structures of CRP in the protein data bank (PDB) that also displayed crystal contacts laterally between pentamers. From the crystal structures available, three formed lateral contacts between CRP pentamers; 3PVN, 3PVO and 7TBA (**Figs. 3, S9 & S10**) (5, 16). In particular, the CRP crystal 3PVN contained dimers of pentamers that were separated by 10 nm and appeared to also fit the density and tomograms (**Figs. 3A & S11**). This model revealed potential interactions between Lys57 and Glu130 on the corner of one CRP pentamer interacting with Glu197 and Lys123 on the edge of an adjacent CRP pentamer, respectively. An equivalent constellation of atoms and interactions is present between crystallographic unit cells in 7TBA (5). Fitting two of these dimers into the map produced a tetrameric platform (**Fig. 3B**). Although no interactions were immediately apparent between the non-crystallographic dimers, Lys57 and Glu130 were again found in close proximity between CRP pentamers, indicating that a pair of salt bridges may stabilise this tetrameric conformation (**Fig. 3B**). Interestingly, Lys57 has been previously shown to be important for CRP-mediated complement activation, an observation that has hitherto been unexplained (44).

**Figure 3.**
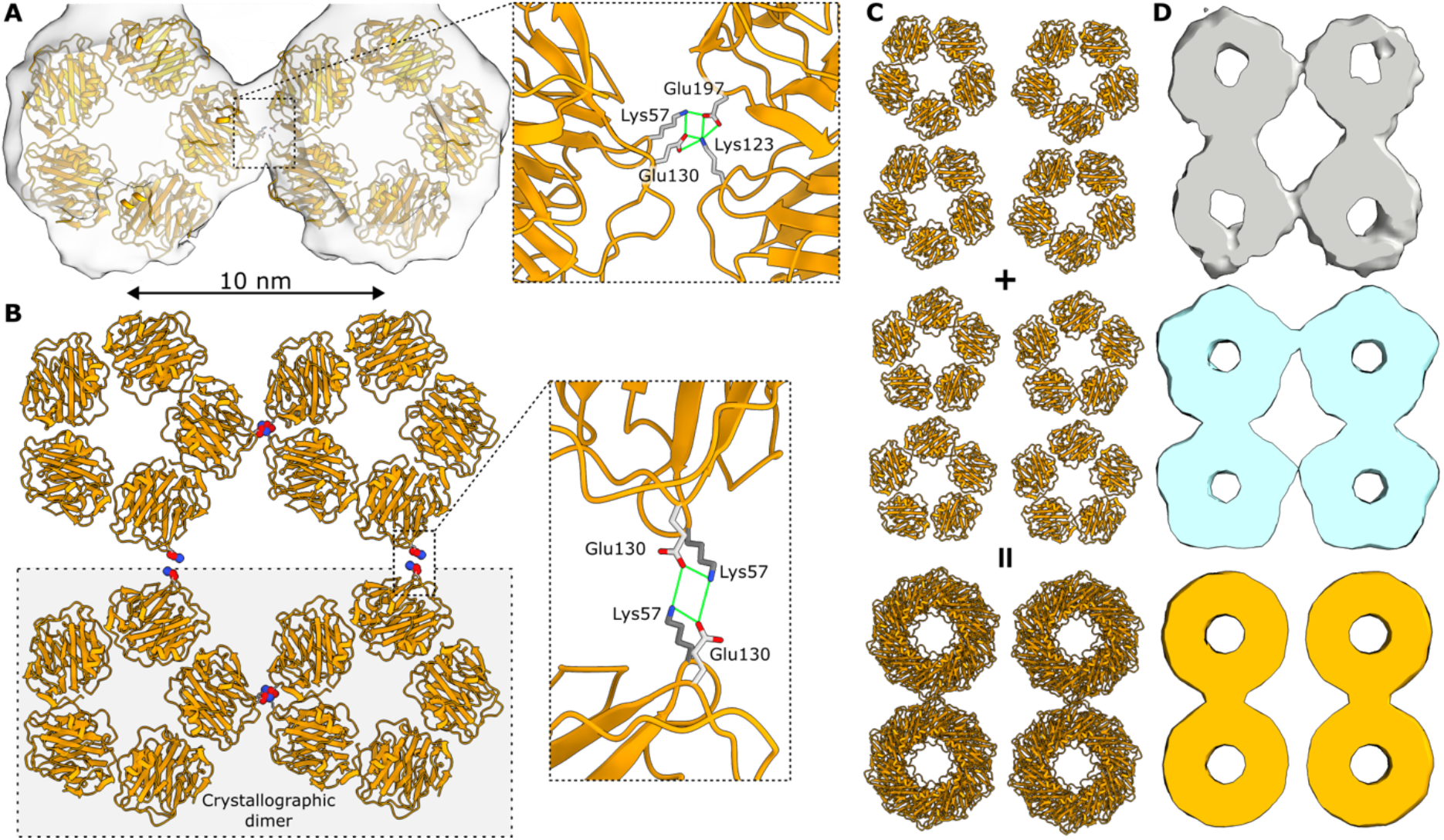
Lateral interactions stabilise tetrameric CRP platforms. **A)** Crystallographic dimer (3PVN) of CRP pentamers (orange) fit into the subtomogram map. Inset shows the existing salt bridge interactions (green lines) between neighbouring CRP complexes. **B)** Tetrameric CRP platform formed from two crystallographic dimers. Inset shows the putative salt bridge interactions (green lines) stabilising the tetramer. **C)** Schematic showing how a superposition of tetramers rotated 180° with respect to each other results in a toroidal appearance. **D)** Maps lowpass filtered to 20 Å: Grey, slice through the subtomogram density; blue, slice through a simulated map of one of the CRP tetramer orientations; orange, slice through a simulated map of the superposed tetramers.

An alternative dimeric configuration found within the crystal with PDB code 3PVO instead formed contacts between two edges of CRP pentamers (**Fig. S10**). This dimer was closer together, separated by 9 nm, and consequently appeared to fit the density less well. Nevertheless, this also exhibited possible inter-pentamer contacts between Lys57, Glu130, Lys123 and Glu197 within the crystallographic dimer, and Lys57 and Glu130 were again adjacent to one another between non-crystallographic dimers (**Fig. S10**). Dimers of CRP adopting this closer configuration were also apparent in tomographic slices (**Fig. S11**).

The toroidal maps of CRP may be explained by the subtomogram average being a superposition of CRP tetramers in orientations that are rotated 180° with respect to each other (**Fig. 3C**,**D**). Since multireference refinement could not separate these two populations (**Fig. S8**), these are presumably equally populated and able to recruit the C1 complex.

### Hexameric C1 binds to tetrameric CRP platforms

Four of the gC1q domains bound within the hole of the toroidal CRP pentamers (**Fig. 2C**,**D**), and the remaining two gC1q domains hover in a defined location ∼6.5 nm above the surface (**Fig. 2D**). The gC1q domain is too large to fully enter the hole in the ring of pentameric CRP (**Fig. S12A**). Instead, the gC1q domains bound in an off-centre manner, preferentially contacting the inner edge of the cavity furthest from the centre of the CRP-C1 complex (**Figs. 4A & S12B**). Although the tetrameric CRP platform forms a 10×11 nm rectangle, this off-centre binding means that the centres of the gC1q domains are placed on a 12×13 nm rectangle, which differs from previous hexagonal arrangements by antibodies, where gC1q heads were separated by ∼12 nm (**Fig. 4B**).

**Figure 4.**
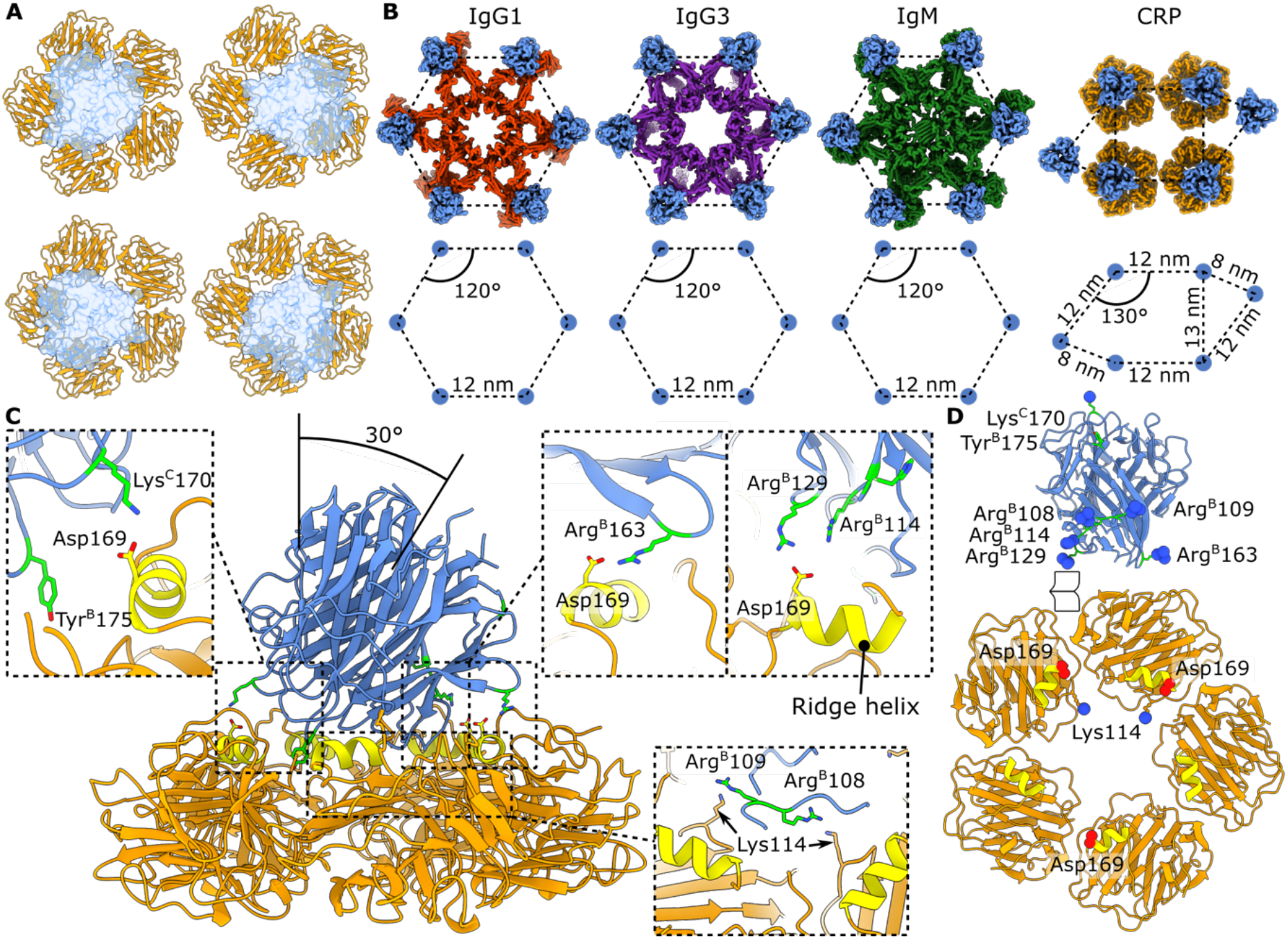
C1q binds to tetrameric CRP platforms. **A)** Binding location of the four gC1q domains on the tetrameric CRP platform. **B)** Measured distances between gC1q domains bound to IgG1, IgG3, IgM and CRP. **C)** Interactions suggested by our model between pentameric CRP and gC1q. The CRP ridge helix residues are coloured yellow, and interacting residues on gC1q are coloured green. **D)** Unfolding the interface between gC1q and CRP to visualize the interacting residues. Oxygen and nitrogen atoms involved in salt bridges are shown as red and blue spheres, respectively.

The resolution of the map precluded accurate determination of the orientation of the gC1q domains, although density connecting the gC1q domains to the C1r_2_s_2_ protease platform suggested that they were tilted by 30° from perpendicular to the membrane (**Fig. 4C**). Residues Pro168-Gly177 on the A-face α-helix of CRP, known as the ridge helix, have been shown to be important in C1 binding (45). Previous mutagenesis studies on purified gC1qA, gC1qB and gC1qC chains have suggested two patches of residues on the gC1q domain that may impact gC1q binding to CRP (29). These are located at the apex of headpiece (Lys^C^170 and Tyr^B^175 on C1qC and C1qB, respectively) and on the edge of C1qB (Arg^B^114, His^B^117, Lys^B^136 & Arg^B^163, **Figs. 4C & S12C**). The gC1q domains were placed in the map manually to account for these interactions, before the fit was optimised within ChimeraX (46) and allowed to relax by briefly simulating within ISOLDE (47).

The fitted and relaxed model shows several interactions of note between gC1q and CRP. In particular, gC1q residues Arg^B^114, Arg^B^163, and Lys^C^170 all interact with Asp169 from three different CRP monomers (**Fig. 4C**,**D**). Although Arg^B^129 has not been previously shown to be important for C1 binding to CRP, it has been shown to be important for C1q binding to other ligands (6), including IgG1 (33). Here, Arg^B^129 is found interacting with the same Asp169 on CRP as Arg^B^114. Mutation of the CRP residue Asp169 to an alanine has been shown to reduce C1 binding, but the conservative mutation to glutamate did not affect binding (45), indicating a salt bridge interaction with C1q as observed here. Interestingly, mutation of the positively charged Lys114 from CRP actually increased C1q binding and C3 deposition (44, 45), indicating a non-favourable charge interaction with gC1q. Here, we observe Lys114 in close proximity to Arg^B^108 and Arg^B^109, which would repel one another, offering an explanation for the increased binding when removing the Lys114 positive charge. Arg^B^108/109 is known to be important for C1q binding to the long pentraxin PTX3 and IgM (30), suggesting they may also be involved in interactions with CRP. Tyr^B^175 is within ∼6 Å of the CRP ridge helix and therefore may form interactions that are not apparent in this model. Neither His^B^117 or Lys^B^136 are proximal to CRP in this model, and so their previously determined impact on binding and complement activation may be due to other factors (29). Whilst this model provides a structural explanation for previous biochemical data (29, 44, 45), the resolution of the map limits interpretation. Therefore, other orientations and interactions are possible, and we cannot exclude further structural rearrangements that may occur upon C1 binding and activation.

### CRP-bound C1 adopts a pre-C4 cleavage conformation

As observed in the raw tomograms (**Figs. 2A & S6**), the C1r_2_s_2_ protease platform, comprising complement C1r/C1s, Uegf, Bmp1 (CUB)1, epidermal growth factor (EGF) and CUB2 domains (**Fig. S1**), was parallel to the membrane (**Figs. 2B**,**D & S6**), in a similar arrangement as observed for C1 bound to IgG1, IgG3 and IgM (33-35). The two C1r serine protease arms, comprising complement control protein (CCP)1, CCP2 and serine protease (SP) domains (**Fig. S1**), were horizontal, oriented parallel to the liposome surface in the same conformation as previous structures of C1 (34, 35). The C1s arms were oriented slightly downwards, ∼15° from horizontal (**Fig. 2B**,**D**), with the C1s SP domains only 5 nm away from the C1r SP domains (**Fig. 5A**). Although both C1r arms were well resolved, one of the C1s arms was less well resolved than the other (**Fig. 5A**). Furthermore, we observed weak density protruding from the side of the C1r_2_s_2_ platform that could correspond to the C1s CUB2 domain adopting an alternate conformation (**Figs. 5B & S13**). Focussed classification and refinement around one of the protruding densities did not resolve the full C1s arm, but did extend the density somewhat to include the CUB2 and CCP1 domains, allowing a more accurate placement of the complete C1s arm (**Figs. 5C & S13**). In this conformation, the serine protease domains of C1r and C1s were located ∼20 nm apart. This alternate conformation was similar to previous structures of C1 bound to antibody platforms (34, 35), which showed the C1s arms bending down towards the liposome surface where it interacted with C4b, one of the products of the C1s enzyme. In those instances, C1s was found either 5 nm away from C1r when not interacting with C4b, or 16 nm away from C1r when bound to C4b (**Fig. 5C**).

**Figure 5.**
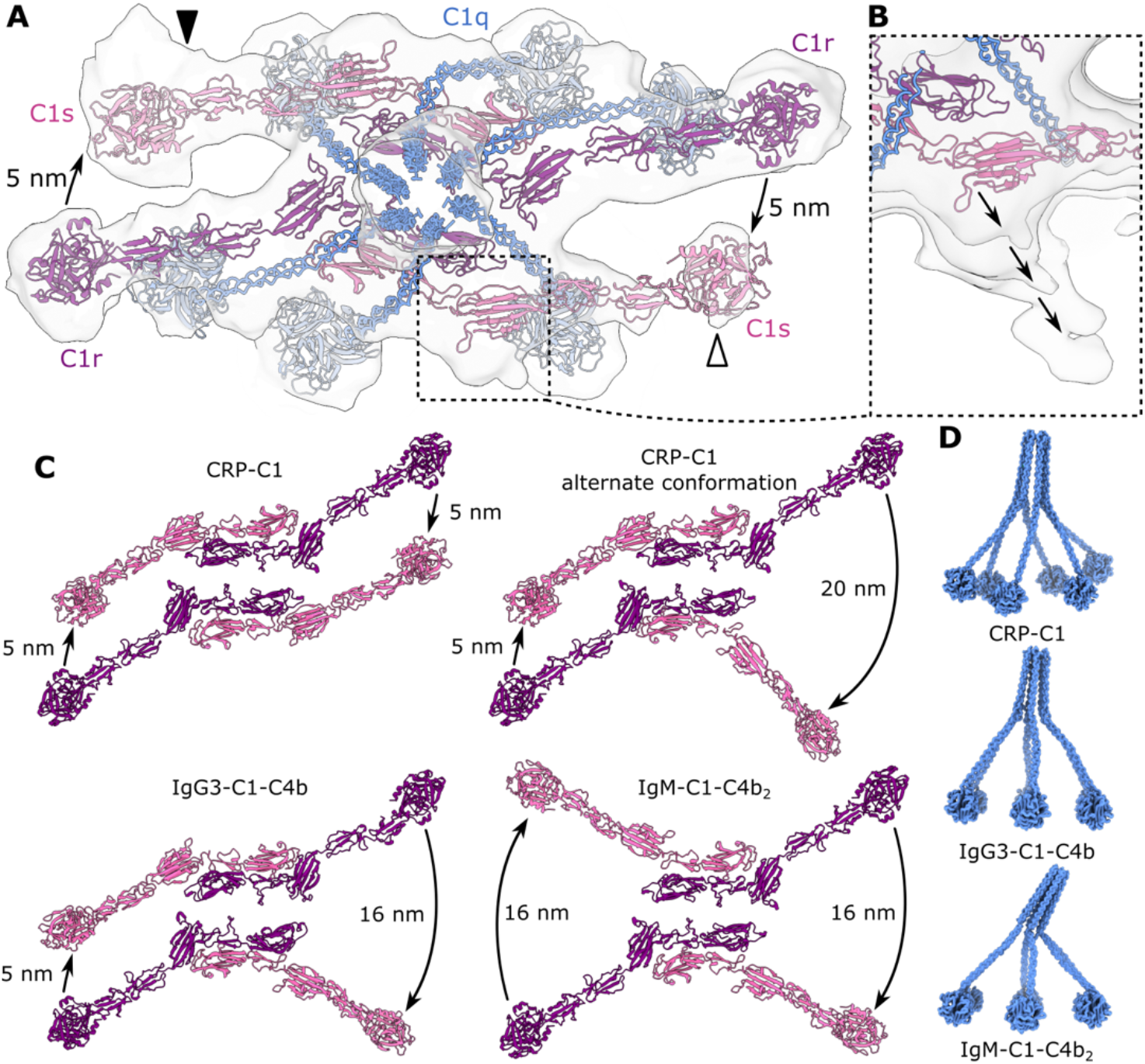
C1 proteases structures indicate a via large domain shift during autoactivation. (**A**) Structure of the C1r2s2 protease platform in PC-CRP-C1. One C1s arm is less well resolved than the other (white and black arrowheads, respectively). (**B**) Focussed classification and a low isosurface threshold indicate extra density at the C1s arm region (black arrows) indicate an alternative conformation. (**C**) Models of C1r (purple) and C1s (pink) from C1 complexed with CRP (focused refinement & normal refinement), IgG3 and IgM (EMDB codes; EMD-18830, CRP; EMD-16251, IgG3; EMD-4878, IgM). Distances between opposing C1r and C1s arms are shown. (**D**) The C1q stalk is upright on IgG3 and CRP, but tilts on top of IgM.

These data suggest that the C1 proteases predominantly adopt a conformation that separates the SP domains by only 5 nm when bound to tetrameric CRP platforms. We have previously observed such a conformation in maps of C1 bound to IgG3 (34), and posit that this close association of the C1r SP to the C1s SP may represent the conformation that C1s adopts before cleavage and activation by the C1r SP. This may also be the ‘resting-state’ of C1, before C1s binds and cleaves C4 and deposits C4b on the surrounding surface. Additionally, the stalk appeared upright in a similar manner to IgG3-C1, but different to IgM-C1 (**Fig. 5D**), where the stalk domain was tilted ∼20° towards the deposited C4b molecule (35). This may represent a configuration of the stalk domain prior to C4 cleavage, as has been hypothesised previously (34).

## Discussion

Despite abundant evidence demonstrating the importance of CRP for human health (1, 12, 48, 49), CRP is underrepresented as a target for therapeutic intervention (50). This absence of therapeutics is a result of the lack of structural information regarding the pentraxin family, which limits our understanding of how they function. We have recently used cryoEM to image solution-phase CRP, which showed the presence of both pentameric and decameric complexes (15). When we attempted to image ligand-bound CRP using cryoET, however, we observed aggregation on the grid (**Fig. S5**), which we discovered was caused by CRP-mediated liposome agglutination (**Fig. 1A**). To achieve a thin sample of ligand-bound CRP complexes, a mutant of CRP (R188A) that is deficient in decamer formation was expressed and purified, leading to significantly less CRP induced agglutination of liposomes (**Figs. 1D & S5**). Whilst this allowed us to proceed with cryoET, the aggregation of lipidic structures observed here with wild-type CRP likely represents a genuine biological function of CRP *in vivo*. Antibodies aggregate antigen-containing membranes in a process of agglutination (51, 52), mediated via the F(ab’)_2_ fragment of antibodies (51), to facilitate the clearance of antigens *in vivo*. Similarly, agglutination of lipid membranes by CRP has been hypothesised to be part of the ability of CRP to help facilitate clearance of antigens by a similar mechanism. Indeed, CRP has been shown to facilitate phagocytosis in various experimental models (20, 53-55). This suggests that decameric CRP is the functional homolog of the F(ab’)_2_ antibody fragment in agglutination and phagocytosis.

Biophysical and cryoEM analysis of liposomes containing LPC, the lipid found on apoptotic cell blebs (17) and bacteria (56) to which CRP binds *in vivo* (17), revealed small, unstable lipid droplets and liposomes that were unsuitable for structural analysis (**Fig. S2**). Instead, we synthesised a hapten-like conjugate comprising a PC headgroup linked to an azide moiety (**Supplementary methods**). We were able to incorporate a controlled concentration of DBCO-modified lipids into known stable liposome formulations during production(57), before copper-free click ligation of the azide-PC molecule with the DBCO-coated liposomes. This resulted in a native CRP ligand stably displayed on liposome surfaces, affording control over sample preparation for cryoET. Combined with the R188A CRP point mutant described above, this resulted in ideal thin samples for cryoET and subtomogram averaging (**Figs. 2 & S5**).

Analysis of ligand-bound CRP via cryoET revealed that CRP associates laterally to form tetrameric arrays on membranes displaying PC (**Fig. 2**). The subtomogram map of CRP tetramers enabled a model to be built, constructed from crystallographic dimers-of-pentamers fit into the map (**Fig. 3**). Laterally-associated dimers were found in three crystal structures of CRP; 3PVO, 3PVN and 7TBA (5, 16) (**Figs. 3, S9 & S10**), and dimers of CRP pentamers were also observed on liposomes in the absence of C1 (**Fig. S11**), suggesting they precede C1 binding and complement activation. Moreover, three PDB entries relating to the other human short pentraxin, serum amyloid component (SAP; 1SAC (58), 4AYU (59) and 3KQR (60)), also contained similar lateral arrangements of pentamers (**Fig. S14**). These structures, together with our cryoET map, indicate that formation of lateral contacts between membrane-bound short pentraxins may lead to tetramers or even higher-order 2D arrays. Structural analysis of the resulting CRP tetramers indicated several residues likely to be important for tetramer formation (**Fig. 3**). These included Lys57, which was shown to be important for complement activation from mutagenesis studies (44). However, Lys57 is far from the postulated C1 binding site (45), and so the reason for this importance was not previously apparent. Here, we provide a structural explanation; CRP must associate via lateral interactions to enable C1 to bind to the A-face. Complement activation therefore requires two sites on CRP; one for lateral association and one for C1 binding, and Lys57 belongs to the former.

A-face residues important for C1 binding and activation have been identified by mutagenesis (44, 45), along with two patches of residues on gC1q that have been shown to be important for CRP binding (29) (**Fig. S12C**). Although the resolution of our map is too low to determine the specific interactions between CRP and the gC1q headgroups, taking these residues into account enabled placement of the gC1g domains into the density, whilst also ensuring the N-termini of each gC1q was correctly oriented towards the density connecting to the C1r_2_s_2_ platform (**Figs. 2 & 4**). This model indicated several salt bridges between Asp169 on CRP with residues Arg^B^114, Arg^B^129 and Arg^B^163 from C1qB, and Lys^C^170 from C1qC (**Fig. 4**). Several of these residues have been previously implicated in CRP-mediated complement activation, but our structural data also provide more sites for future site-directed mutagenesis. Furthermore, Lys114 on CRP was shown to be detrimental for CRP-mediated complement activation, which we now show is likely due to repulsive interactions with Arg^B^108 and Arg^B^109. These repulsive interactions may potentially limit solution-phase C1 binding until CRP tetramerization occurs, leading to higher avidity binding that overcomes this repulsion. Lys114 is non-conserved amongst human pentraxins, and is replaced by a glutamic acid in PTX3, possibly forming an additional salt bridge with gC1q. Whether this leads to relatively enhanced complement activity remains to be seen.

C1 can be activated by hexameric or pseudo-hexameric antibody complexes (33-35, 40), and it was unknown if a similar mechanism existed for pentraxin-mediated complement activation (28, 61). Although hexameric antibody platforms produce maximal complement activation, C1 activation has been shown to require at least two antibodies (62), and tetramers have been shown to be less effective than hexamers (63). Here, we provide the first structure of a pentraxin array binding to the C1 complex and activating the complement system (**Fig. 2**), which revealed C1 binding to tetrameric CRP platforms. Laterally interacting CRP pentamers are therefore likely the functional equivalent of hexameric antibody platforms that are requisite for C1 binding and complement activation (33-35, 40, 64) (**Fig. 4B**).

CRP is known to result in less terminal pathway activation and reduced MAC pore formation than antibody-mediated activation, even with robust C1 activation and C4 cleavage, representing its role in the silent clearance of apoptotic cells (36, 37). Whilst this effect is linked to associations with regulatory proteins such as factor H (37), this does not preclude structural factors playing a role. For instance, tetrameric arrangements of CRP may activate the C1 complex in a less efficient manner than hexameric antibody platforms, resulting in less MAC formation. Indeed, direct comparisons of CRP- and antibody-mediated complement activation shows that CRP results in markedly less MAC mediated lysis (36, 63). Alternatively, formation of an extended 2D array of laterally-associated CRP pentamers could impede deposition of complement components, limiting convertase formation and MAC pore formation. The overall height of the CRP-C1 complex may also influence complement activation. Previously determined structures of C1 bound to antibody platforms have revealed deposited C4b adjacent to or on the antibody complexes themselves (34, 35). IgG1-C1, IgM-C1 and IgG3-C1 complexes are 35, 35 and 49 nm tall, respectively (33-35), and are all tall enough to accommodate C4b (**Fig. S15A**). In contrast, the subtomogram map presented here shows that the CRP-C1 complex stands 29 nm tall. Aligning C1s with the homologous structure of MASP2 bound to C4 (65) illustrates how the height of CRP-C1 does not allow deposition of C4b in the same configuration as for antibody-C1 complexes (**Fig. S15B**). Instead, C4 cleavage by CRP-C1 must occur in a conformation not yet observed. Deposition of C4b is followed by C2 binding, forming the C4bC2 proconvertase, before cleavage of C2 by C1s to form the C4b2b C3 convertase (32). The limited height of CRP-C1 complexes will make cleavage of C4bC2 unlikely. Instead, it may become more likely that C4b is deactivated by Factor I (66), forming the inactive C4d side product. These geometric limitations of the short CRP-C1 complex may also contribute to the reduced complement activating abilities of CRP compared to the taller antibody complexes, as CRP-C1 is less able to form C3 convertases and propagate the complement pathway.

Although the gC1q domains adopted different configurations on the tetrameric CRP array compared to hexameric antibody Fc platforms (**Fig. 4B**), the C1 proteases were highly similar to previous structures (**Figs. 5A & S15**) (33-35). The C1r_2_s_2_ protease platform comprises a dimer of C1rs dimers, where the interface between the two dimers is formed from the C1r CUB1-EGF-CUB2 domains (**Fig. S1**) separated by two collagenous arms of C1q. Opening of this interface was previously posited to precede C1 autoactivation (35), and the similar protease conformation observed in the CRP-C1 structure here indicates a shared activation mechanism with antibody-mediated activation. The protease arms of C1s have also been shown to be highly flexible in other structures of C1 (34, 35), due to the lack of interaction between lysine residue Lys^A^59 of C1qA and the Ca^2+^ ions chelated within the C1s CUB2 domains(67). This weak binding explains the high flexibility of the C1s protease arms around C1q, which appears necessary for C1s to adopt a conformation close to C1r for autoactivation, but still able to rotate around the EGF/CUB2 interface to deposit C4b onto the surrounding surface (35). Indeed, although the major population of CRP-C1 particles showed the SP domains of C1r and C1s physically close to one another in space (**Fig. 4A**), a subpopulation of particles indicated a different orientation of C1s, instead oriented down towards the liposome membrane and ∼20 nm away from C1r (**Figs. 5C & S13**). This orientation was similar to those found in maps of IgM-C1 and IgG3-C1 complexes (34, 35), where C1s was observed bound to deposited C4b. Here, samples were prepared in the absence of C4, which is in contrast to IgM-C1 and IgG3-C1 maps that were imaged in the presence of human serum containing C4 (34, 35). This lack of C4, the substrate for C1s, may be the explanation for C1s orientation. In agreement with this notion, IgG1-C1 complexes imaged using purified complement proteins with catalytically-inactive C1r_2_s_2_ domains was also postulated to form a similar arrangement (33), supporting the idea this C1s conformation represents the pre-activation conformation of the C1 complex. Additionally, a similar arrangement was observed for the IgG3-C1 complex, where a superposition of two C1s conformation was resolved (34). The structure of IgG3-C1 was determined in the presence of C4, and therefore also represents the ‘active’ state of the C1 complex. Together, these structures indicate that the resting state of the C1 complex is one where C1s is proximal to the C1r proteases, either before activation by C1r, or before cleavage of C4.

In conclusion, by combining a synthetic CRP ligand with structure-guided mutagenesis that decoupled complement activation from agglutination, we were able to form ligand-bound CRP-C1 complexes ideal for cryoEM imaging and provide the first structural data regarding pentraxin-mediated complement activation. These data explain previous, sometimes contradictory, biochemical data regarding C1 binding sites and complement activation by CRP, and provide more insights into future sites for mutation. CRP is a target for therapeutic intervention (5, 68), and the structure of this immunologically pivotal complex presented herein may provide future routes for targeting CRP for therapeutic purposes.

## Methods

### Liposome production

Liposomes were prepared using 1,2-dimyristoyl-sn-glycero-3-phosphocholine (DMPC), 1,2-dimyristoyl-sn-glycero-3-phospho-(1’-rac-glycerol) (DMPG), cholesterol (chol), 1,2-dioleoyl-sn-glycero-3-phosphoethanolamine-N-dibenzocyclooctyl (DBCO) or 1-myristoyl-2-hydroxy-sn-glycero-3-phosphocholine (LPC) purchased from Avanti Polar Lipids (USA). Lipid films were composed of DMPC:chol:DMPG:DBCO (40:50:5:5 mol%), DMPC:LPC (80:20 mol%), LPC (100 mol%) or DMPC:chol:LPC (60:0:20, 50:10:20, 40:20:20 or 30:30:20). Components were dissolved in chloroform:methanol (9:1, v/v) before drying under nitrogen gas and desiccation overnight. Films were rehydrated at 60°C or room temperature, for DBCO containing liposomes and LPC formulations respectively, for 1 hour in 50 mM Tris, 150 mM NaCl, 5 mM CaCl_2_ pH 7.4 (TBS) to a final lipid concentration of 0.8 mg/ml. Then the azide-PC compound was used to modify the DBCO liposomes. Lipids and the PC compound were reacted overnight at a 50:1 azide-PC:DBCO ratio under continuous agitation and kept at RT. Lipids were centrifuged at 21000g and supernatant with excess azide-PC compound was removed before lipids were resuspended into buffer. Hydrated lipid films were then agitated and vortexed for 10 seconds before being sequentially extruded through 400 nm and 100 nm membranes, with 11 passes each, using an extruder from Avanti Polar Lipids (USA).

### Agglutination assay

Agglutination of lipids by WT CRP was monitored in real time on a Clariostar plus plate reader (BMG Labtech, Germany). Liposomes were added to wells of a 96 well plate, before CRP (1000 nM) or TBS buffer control was added. The plate was then inserted in the plate reader and the OD_600_ was measured. To judge the effectiveness of the R188A mutant on reducing agglutination an endpoint assay was used. Briefly, the OD_600_ of liposomes without added protein was measured for two minutes on a Clariostar plus plate reader (BMG Labtech, Germany) before WT or R188A CRP (1000 nM) was added to the liposomes and incubated at room temperature for ∼5 minutes prior to the OD600 being measured again for two minutes. The values were normalised to the highest value in the plate, before the initial baseline was subtracted as background. Values across triplicate wells were then averaged with the standard deviation also being calculated.

### Production CRP mutant R188A

Wild type CRP was ordered from Sigma (Missouri, United States). A DNA construct for CRP with the mutation arginine to alanine at position 188 (R188A) and preceded by a signal peptide for secretion, was codon optimized for expression in human cells and ordered from GeneArt, ThermoFisher Scientific (Regensburg, Germany). In the amino acid sequence below, the signal peptide is indicated in bold, and the R188A mutation is underlined:

> **MEFGLSWVFLVALLRGVQC**QTDMSRKAFVFPKESDTSYVSLKAPLTKPLKAFTV CLHFYTELSSTRGYSIFSYATKRQDNEILIFWSKDIGYSFTVGGSEILFEVPEV TVAPVHICTSWESASGIVEFWVDGKPRVRKSLKKGYTVGAEASIILGQEQDSFG GNFEGSQSLVGDIGNVNMWDFVLSPDEINTIYLGGPFSPNVLNW<u>A</u>ALKYEVQGE VFTKPQLWP

The DNA construct was cloned by HindIII and EcoRI restriction sites into a pcDNA3.3 vector (ThermoFisher Scientific) behind the CMV promotor. Mutant CRP protein was produced by transfecting 25 µg plasmid DNA into Expi293F™ cells using the ExpiFectamine™ 293 Transfection Kit (ThermoFisher Scientific) according to the manufacturer’s instructions. Supernatant was harvested after 6 days, centrifuged and sequentially filtered through 0.45 µm and 0.2 µm filters (Cytiva) to remove cell debris.

Harvested supernatant was diluted 2× in Tris-Hcl 50 mM NaCl 150 mM CaCl2 2 mM pH 8.0 (binding buffer) and incubated with immobilized p-aminophenyl phosphoryl choline beads (ThermoFisher Scientific) at room temperature for one hour. Beads were then pelleted at 5000 gs and the supernatant was discarded, before adding two column volumes of binding buffer and a further 5 minute incubation at room temperature. The beads were then pelleted as before, with the subsequent application of two columns volumes of Tris-HCl 50 mM NaCl 150 mM EDTA 2 mM pH 8.0 (elution buffer) added. The sample was then incubated and pelleted as before. The eluate was then dialysed against Tris-HCl 50 mM, NaCl 150 mM pH 7.4 (TBS) and 2 mM CaCl_2_ overnight, before analysis via SDS-PAGE. Fractions containing CRP were then concentrated in spin filters with a 10 kDa cut-off, before filling the filter with TBS with 5 mM CaCl_2_ and concentrating again before storage at 4°.

### Sample preparation for cryoEM

Liposomes (DMPC:Cholesterol:DMPG:PE-DBCO, 40:50:5:5 mol%) were incubated with 500 nM of RA-CRP at 4°C for one hour. Purified C1 complex (Complement technologies, Texas, United States) was buffer exchanged in spin filters with a 50 kDa cut-off to TBS with 5 mM CaCl_2_, before adding at 1000 nM to the RA-CRP liposome mixture and incubated at room temperature for 30 minutes. Samples were then kept on ice until application to the grid, prior to which protein A-coated 10 nm gold colloids were added. Freshly plasma-cleaned 200 mesh copper grids, with lacey carbon support (Electron Microscopy Sciences), were loaded into a Leica EMGP (Leica Microsystems) and 6 µl sample was applied. Grids were blotted from the back for 1-3 seconds at 4°C and 65% humidity. Grids were then clipped and stored in liquid nitrogen.

### CryoET data collection and reconstruction

Tilt series were collected on a Talos Arctica (ThermoFisher Scientific) operating at 200 kV with a K3 direct electron detector and a Bioquantum energy filter (Gatan) with a slit width of 20 eV. Tilt series were acquired using Tomography 5 (version 5.5.0; ThermoFisher Scientific) in counting mode at 49,000× magnification with a pixel size of 1.74 Å using a dose-symmetric scheme from 0° to +/-57°, in 3° increments. A defocus range of -3 µm to -6 µm was used with tracking before each tilt acquisition. A total dose of 60 e-/Å^2^ was used.

Raw frames were aligned using the “alignframes” command from IMOD (version 4.11) (69). Tomograms were reconstructed after 2× binning of tilt series, CTF estimation and erasure of gold fiducials using IMOD (version 4.11) with weighed back-projection.

### Particle picking and subtomogram averaging

Manual picking of 133 cryotomograms using the e2spt_boxer_old.py command in EMAN2 (version 2.91) (70) produced an initial set of 3,428 particles. Particles were extracted with a box size of 168 pixels (3.48 Å/pixel) after contrast inversion and normalization. An initial model consisting of a lipid bilayer model was filtered to 40 Å and white noise was added in EMAN2 (version 2.91) using e2proc3d.py. All subsequent steps were performed using Dynamo (version 1.1.157) (71) unless otherwise stated and are summarised in **Fig. S7**. Using the initial noisy bilayer model, particles were refined to align all the particles to the lipid membrane. A soft mask created using e2filtertool.py in EMAN2 (70) focusing on the CRP platform and lipid bilayer was used to align particles further with C2 symmetry enforced. Particles then underwent rounds of multireference classification to eliminate particles containing only a lipid membrane. The resulting particle stack of 2518 particles were then split into even and odd halves and underwent multiple rounds of alignment, without symmetry enforced, with an oval mask focusing on both CRP and C1, made in EMAN2 as before (70). A new mask focusing on the C1 protease platform was then produced as above, before refining the particles again. Resolutions were estimated by calculating FSC between even and odd data sets using the EMAN2 command e2proc3d.py, showing resolutions of 43 and 24 Å for unmasked and masked maps, respectively. The final map underwent another round of 3D classification to attempt to resolve heterogeneity in the map (**Fig. S8**). Additionally, a focused refinement of the C1s platform was performed on the final map (**Fig. S13**).

### Model building

Six copies of gC1q were fitted into the map as rigid bodies using 1PK6 (72) as a model. Each gC1q domain was oriented as directed by the density and previously published biochemical data on patches of residues on gC1q that interact with CRP (29). The collagen arms of C1q are composed of six heterotrimeric collagen fibrils, which were modelled using ccbuilder 2.0 (73) using default values for radius and pitch. The region of the collagen arms between the C1r_2_s_2_ protease platform and gC1q domains, comprising residues A58-89, B60-91 and C57-88 (A, B and C refer to the individual chain within the heterotrimeric structure), were fit into the map as rigid fibrils between the relevant gC1q domain and their locations in the protease platform, which bind via known lysine residues (67). Next, the C1q stalk, comprising residues A1-39, B1-41, and C1-38 were fit into the map and known disulfide bonds formed between the C-C and A-B interchain cysteine residues. The C1q stalk and collagen region between the C1r_2_s_2_ platform and gC1q domains are separated by residues A40-57, B42-59 and C39-56, which were oriented in the map to connect these two regions. The C1r_2_s_2_ proteases were modelled based on the crystal structure of the CUB1-EGF-CUB2 6F1C (74). The C1r protease arms, comprising the CCP1-CCP2-SP domains, were based on PDB model 1GPZ (75). The C1s protease arms were modelled based on PDB model 4J1Y (76). The different orientations of C1s were modelled by rotating the CUB2 domain around the C1q collagen arms as described in the main text.

## Acknowledgements

This research was supported by a European Research Council Grant 759517 to THS. We gratefully acknowledge the assistance of Leoni Abendstein with cryoET data collection and analysis.

## Author Contributions

DPN performed cryoEM imaging and reconstructions, as well as biochemical assays data analysis and protein purification. MI and DVF synthesised the PC-azide conjugate for presentation on liposomes. DPN and SMWRH performed C1s proteases assays. MK, JW and TvdV helped with optimisation of the biochemical assays. DJD and LAT performed protein expression. DPN and THS analysed cryoEM data. THS built atomic models and supervised the work. DPN and THS wrote the manuscript. All authors contributed to and approved the manuscript.

## Data Availability Statement

The cryo-EM map of liposome bound tetrameric CRP associated with the C1 complex is deposited in the Electron Microscopy database (EMDB) with accession code EMD-18830. All other study data are included in the main text and SI Appendix.

