## Supplemental Information for "Structural basis for surface activation of the classical complement cascade by the short pentraxin C-reactive protein"

### Supplementary Figures

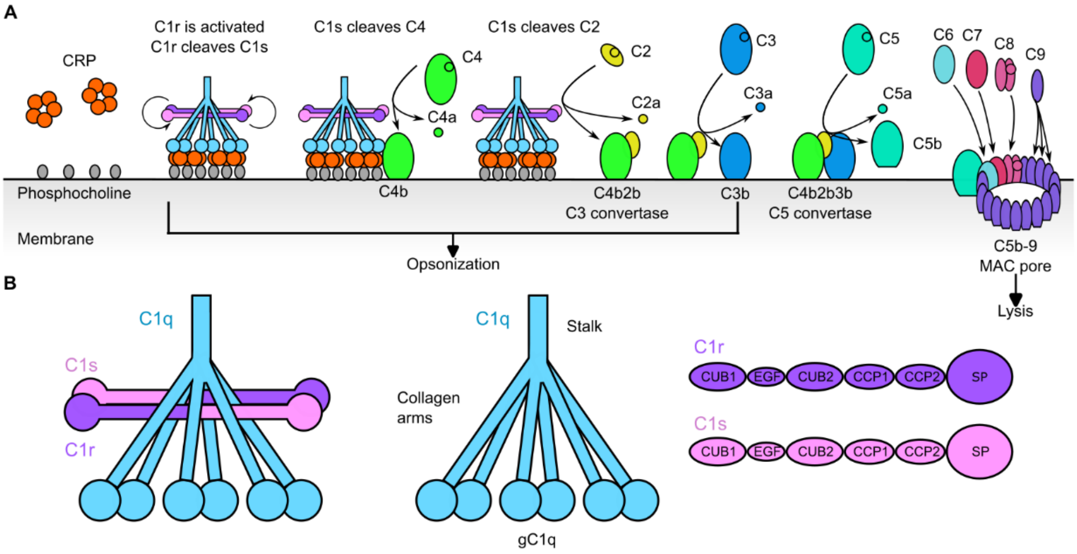


**Figure S1: Schematic of CRP based complement activation and the structure of the C1 complex.** (**A**) Pathway of CRP-mediated complement activation and the physiological outcomes of each step highlighted i.e. opsonization or lysis. (**B**) Schematic of the structure and domain organization of the C1 complex. Abbreviations: CUB, C1r/C1s, Uegf and Bmp1 domain; EGF, epidermal growth factor domain; CCP, complement control protein domain; SP, serine protease domain.

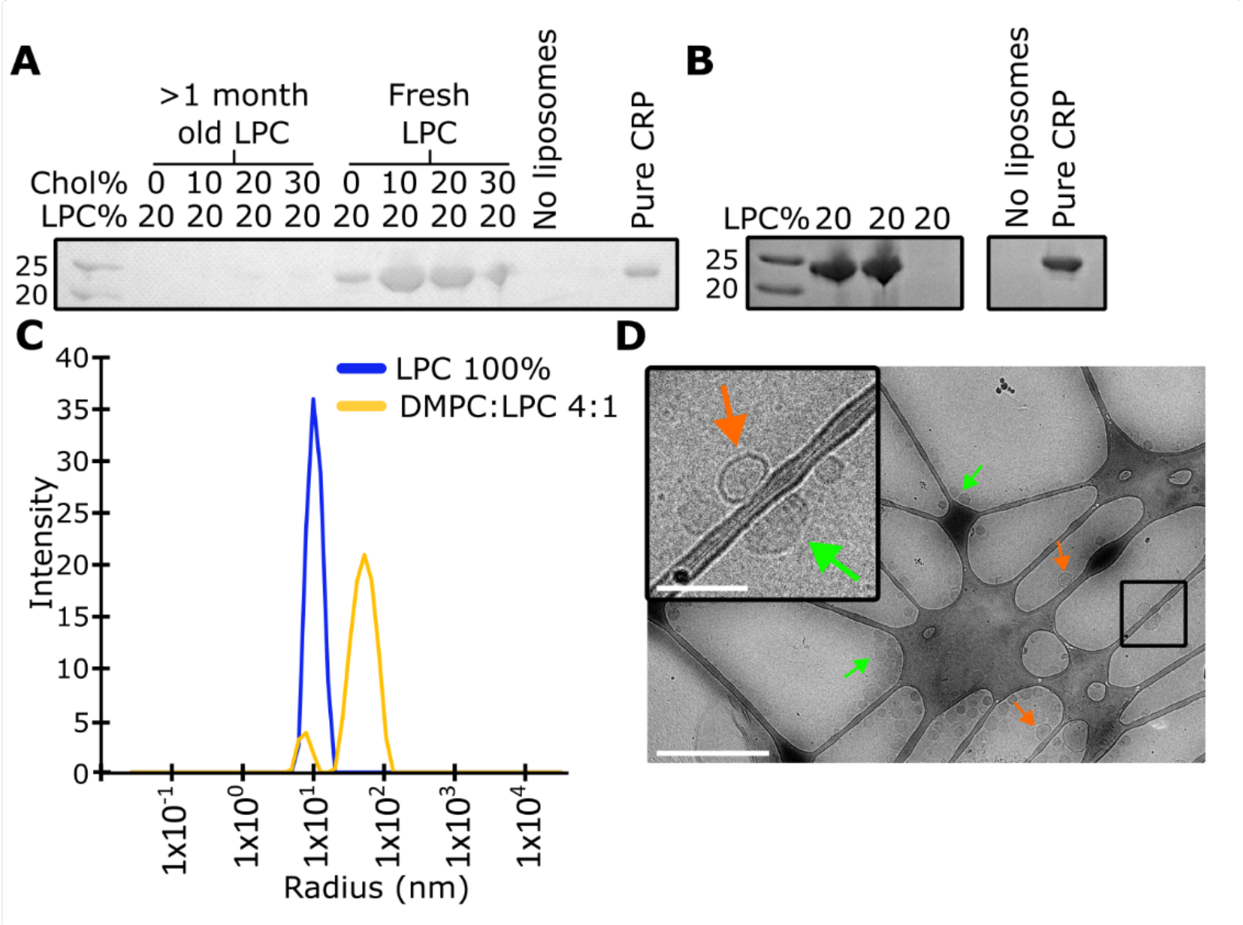


**Figure S2: Liposomes formed from the native CRP ligand LPC are unsuitable for biochemical and cryoEM analysis.** (**A**) Liposome binding assay of liposomes using fresh and old (>1 month) stocks of LPC during liposome synthesis. Cholesterol and LPC content of liposomes is shown (mol%), with the remainder made up by DMPC. (**B**) Liposome binding assay of three liposomes batches containing LPC (DMPC:LPC 80:20% mol%). (**C**) Dynamic light scattering (DLS) of pure LPC and liposomes containing both DMPC and LPC (DMPC:LPC 80:20% mol%). (**D**) Micrograph from cryo electron microscopy analysis of liposomes (DMPC:LPC 80:20% mol%) identifying heterogeneous populations of liposomes on the grid (orange and green arrows indicating two instances of different liposome morphologies). Scale bars represent 500 and 50 nm in the full micrograph and enlarged section, respectively.

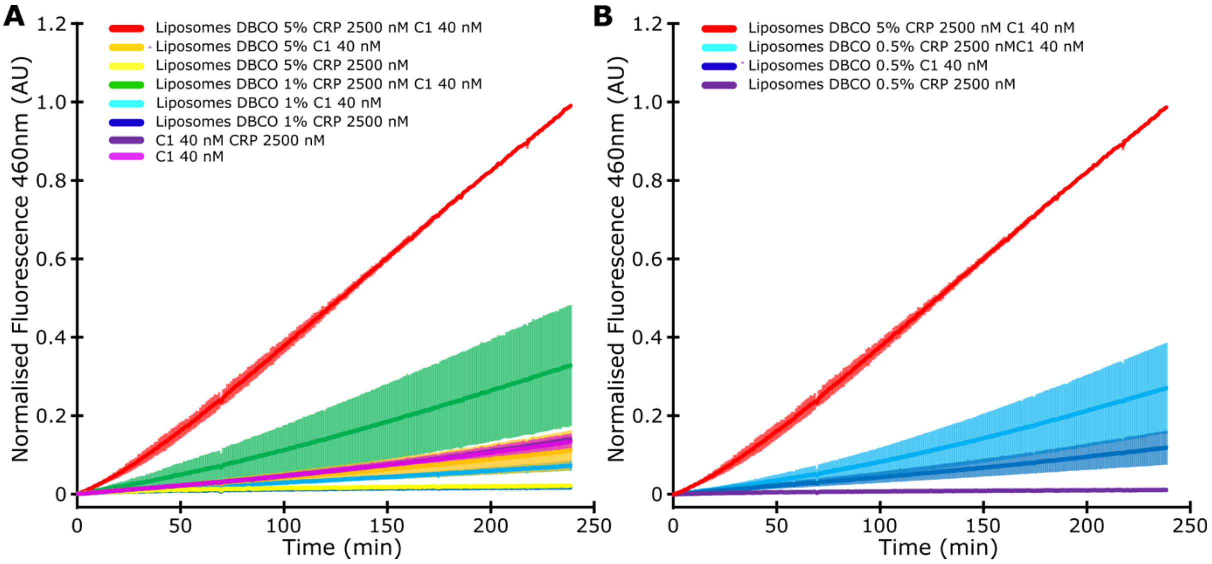


**Figure S3: PC-azide conjugate linked to liposomes via copper free click chemistry binds to CRP to activate the C1 complex.** (**A**) C1s activation assay with 5% and 1% DBCO coated liposomes (DMPC:Cholesterol:DMPG:PE-DBCO, 40:50:5:5 mol%). (**C**) C1s activation assay from the same plates as **B**, showing the 0.5% DBCO liposomes. The 5% DBCO liposomes in the presence of CRP and C1 is shown in both for comparison. Two independent plates were assayed, each containing duplicate wells. The error bars represent the standard deviation between the plates.

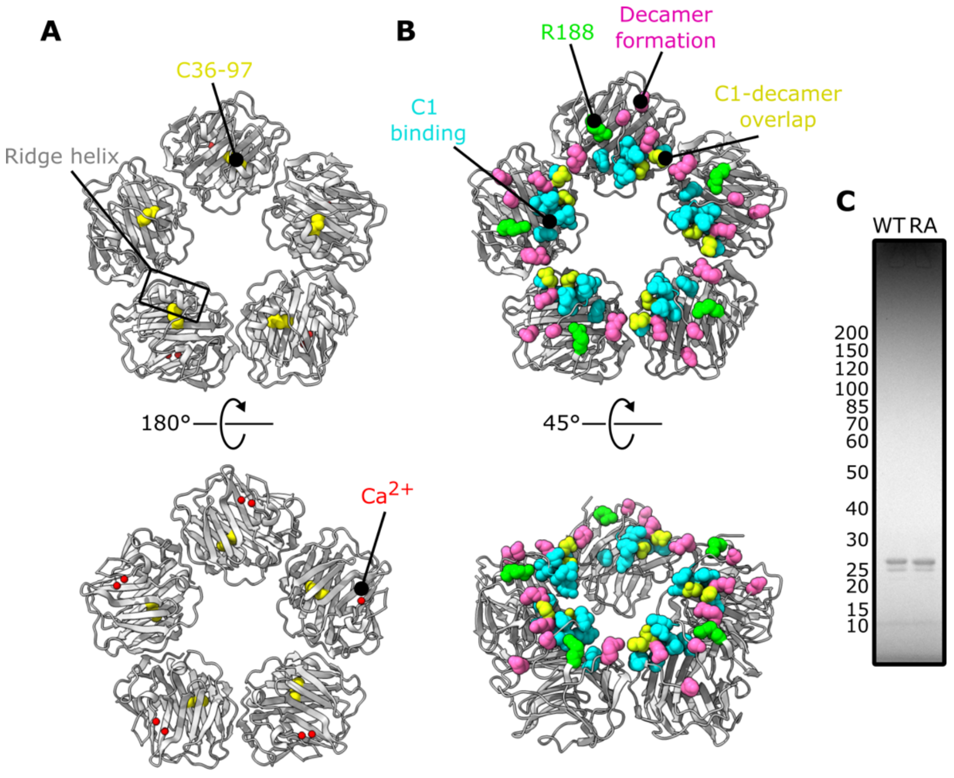


**Figure S4: Structural hotspots on pentameric CRP and biochemical properties of R188A CRP.** (**A**) Location of the disulfide bonds (yellow, C36-C97), $Ca^{2+}$ ions (red) and ridge helix (boxed area). (**B**) Residues on CRP involved in decamerisation (pink, R6, D163, N172 & S181) and C1 binding (cyan, P168-G177 & H38). Residues that are found in both patches (yellow, E170 and D169) are also highlighted. R188, the residues mutated in this study, are shown in green. (**C**) SDS-PAGE analysis of Coomassie stained pure WT and R188A (RA) CRP.

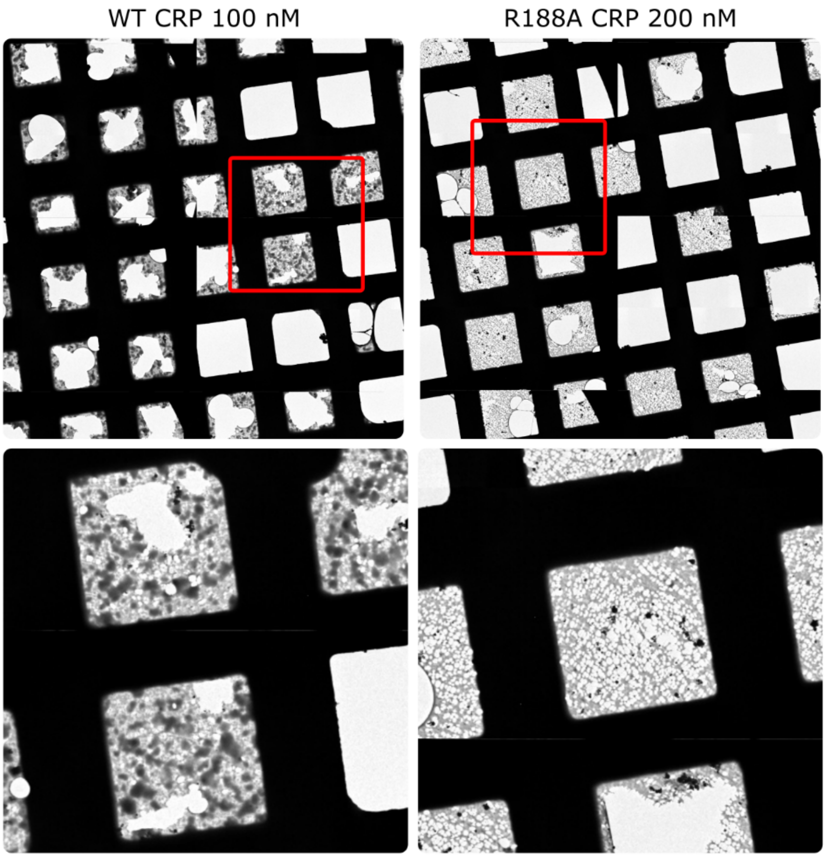


**Figure S5: R188A CRP causes less on grid aggregation than WT CRP.** Representative enlarged imaged from atlas images taken during data collection and screening.

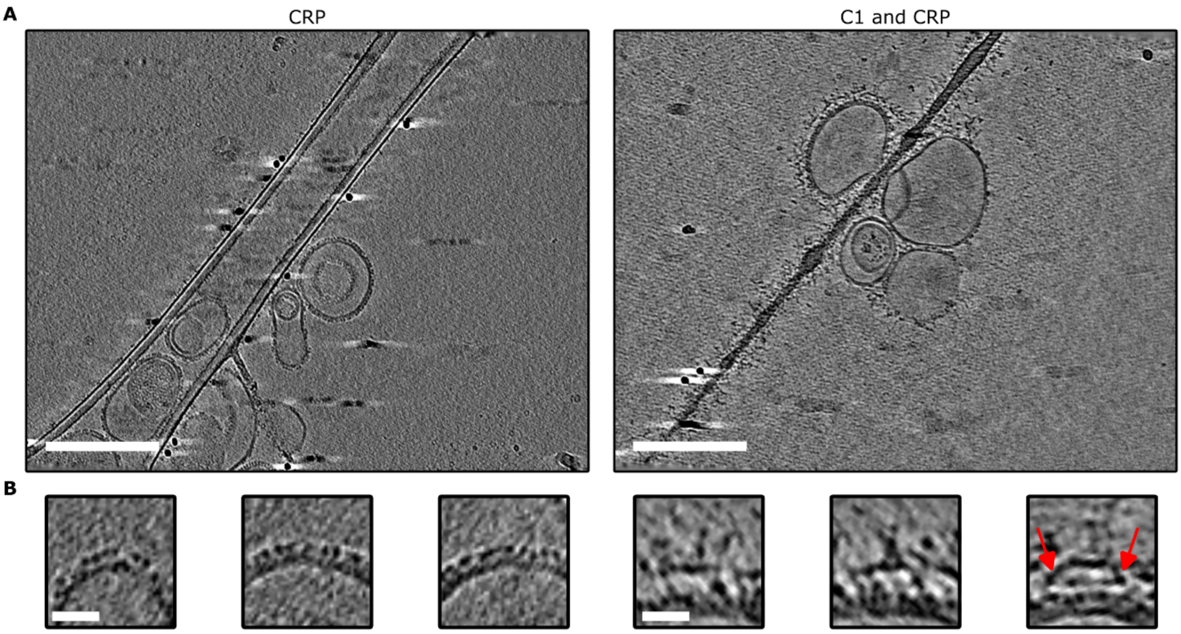


**Figure S6: Slices through cryotomograms of liposomes and enlarged subtomograms of PC-CRP-C1 complexes.** (**A**) Slices 14 nm thick through tomograms of liposomes (DMPC:Cholesterol:DMPG:PE-DBCO, 40:50:5:5 mol%) bound to CRP or C1-CRP. (**B**) Enlarged subtomograms of lipid bound CRP or C1-CRP, with density connecting the CRP layer with the $C1r_2s_2$ protease platform shown by red arrows. Scale bars in **A** and **B** represents 100 and 10 nm respectively.

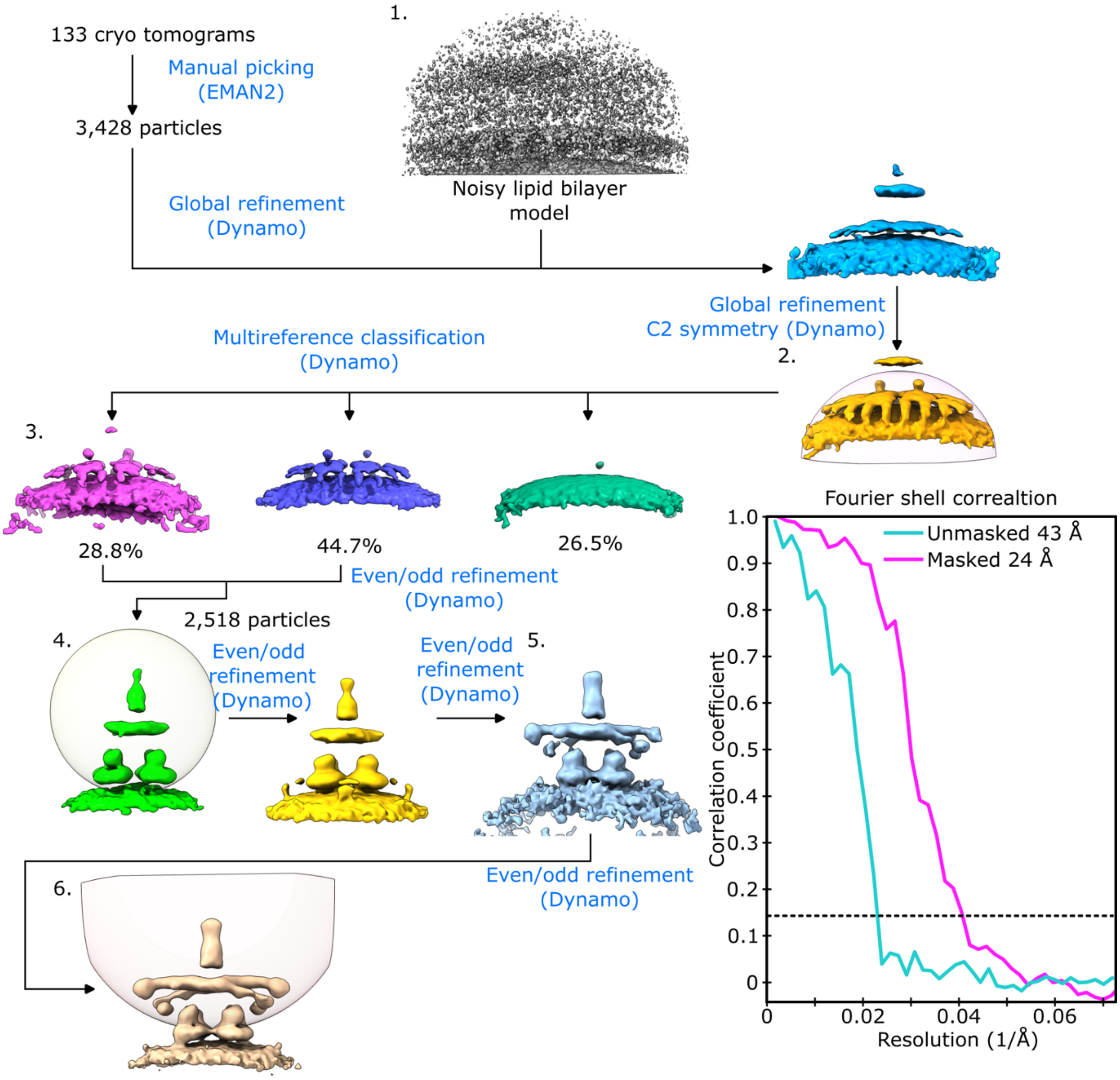


**Figure S7: Subtomogram averaging of PC-CRP-C1 complexes.** A noisy model of a lipid bilayer was used to globally align particles to the membrane (1). These particles were then refined with C2 symmetry enforced and a mask focusing on the CRP platform (2). No further symmetry was used or applied. Classification was then used to remove particles containing only lipid bilayer (3). The remaining 2,518 particles were then split into even and odd data sets and refined with a spherical mask encompassing C1 and the CRP platform (4). This was repeated iteratively two more times with the same spherical mask, using the previous refinements result as an initial model (5). The particles were then refined with a mask focusing on the C1 complex (6). Masks are shown in their first instance**.** The Fourier shell correlation (FSC) value represents the masked and unmasked resolution estimates (FSC = 0.143 cutoff) of the final refinement (6).

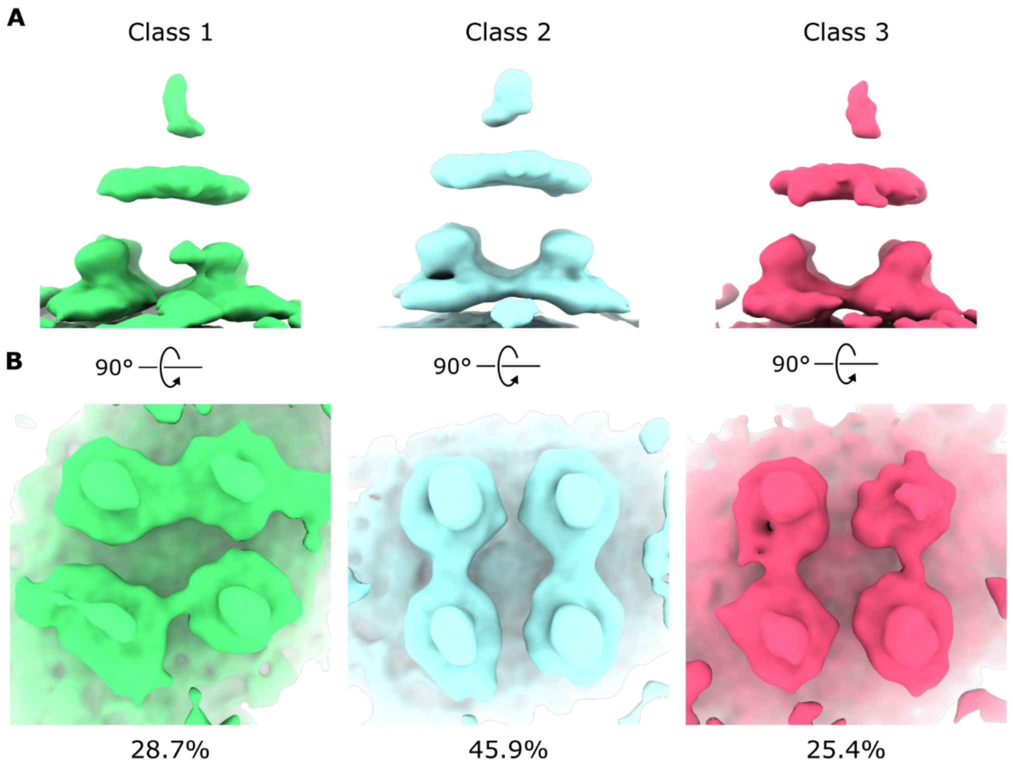


**Figure S8: 3D Class averaging of the final PC-CRP-C1 subtomogram map.** (**A**) 3D class averages showing the entire complex including the C1 stalk and protease platform. (**B**) Subtomogram maps of each class focusing on the tetrameric CRP platform. Proportions of the total number of particles contributing to each class are shown as a percentage below each image.

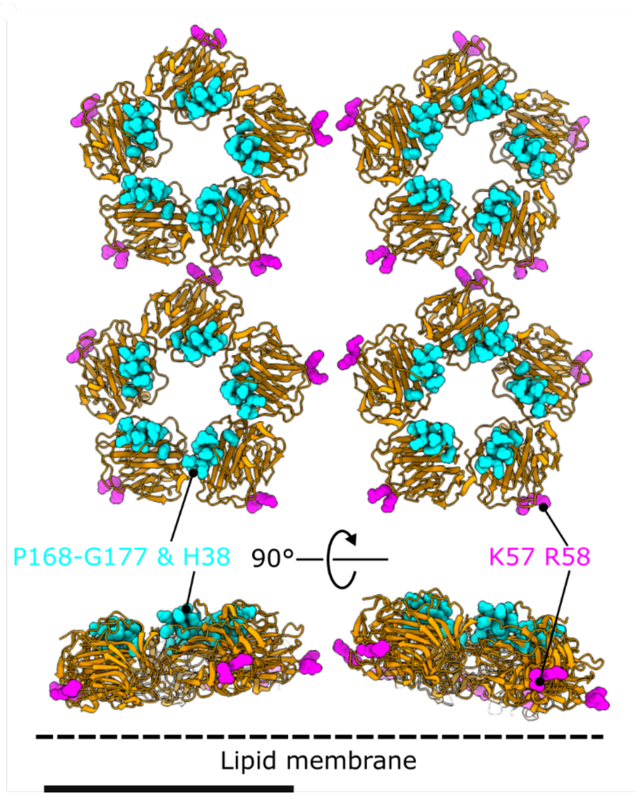


**Figure S9: Locations of residues important in C1 binding and complement activation in the tetrameric platform of CRP.** Tetrameric CRP platforms made up of crystallographic dimers (3PVN) of CRP pentamers (orange), fitted into the subtomogram map. Top and bottom panels show top down and side views respectively, with C1 binding residues (P168-G177 and H38) and residues involved in complement activation (K57 and R58) shown in cyan and purple, respectively. The side facing the lipid membrane is shown by a dashed line in the bottom panel. The scale bar represents 10 nm.

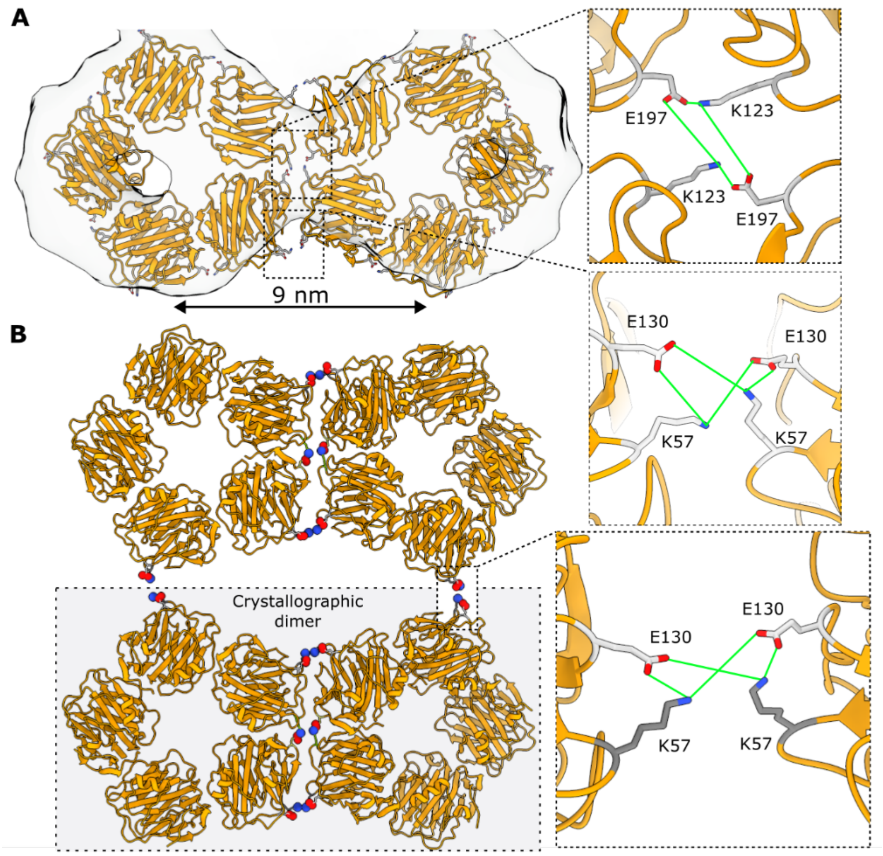


**Figure S10: The crystallographic dimer 3PV0 fit into the subtomogram map. A)** Crystallographic dimer (PDB code 3PV0) of CRP pentamers (orange) fit into the subtomogram map. Inset shows the potential salt bridge interactions (green lines) between neighbouring CRP complexes. **B)** Tetrameric CRP platform formed from two crystallographic dimers. Inset shows the putative salt bridge interactions (green lines) stabilising the tetramer.

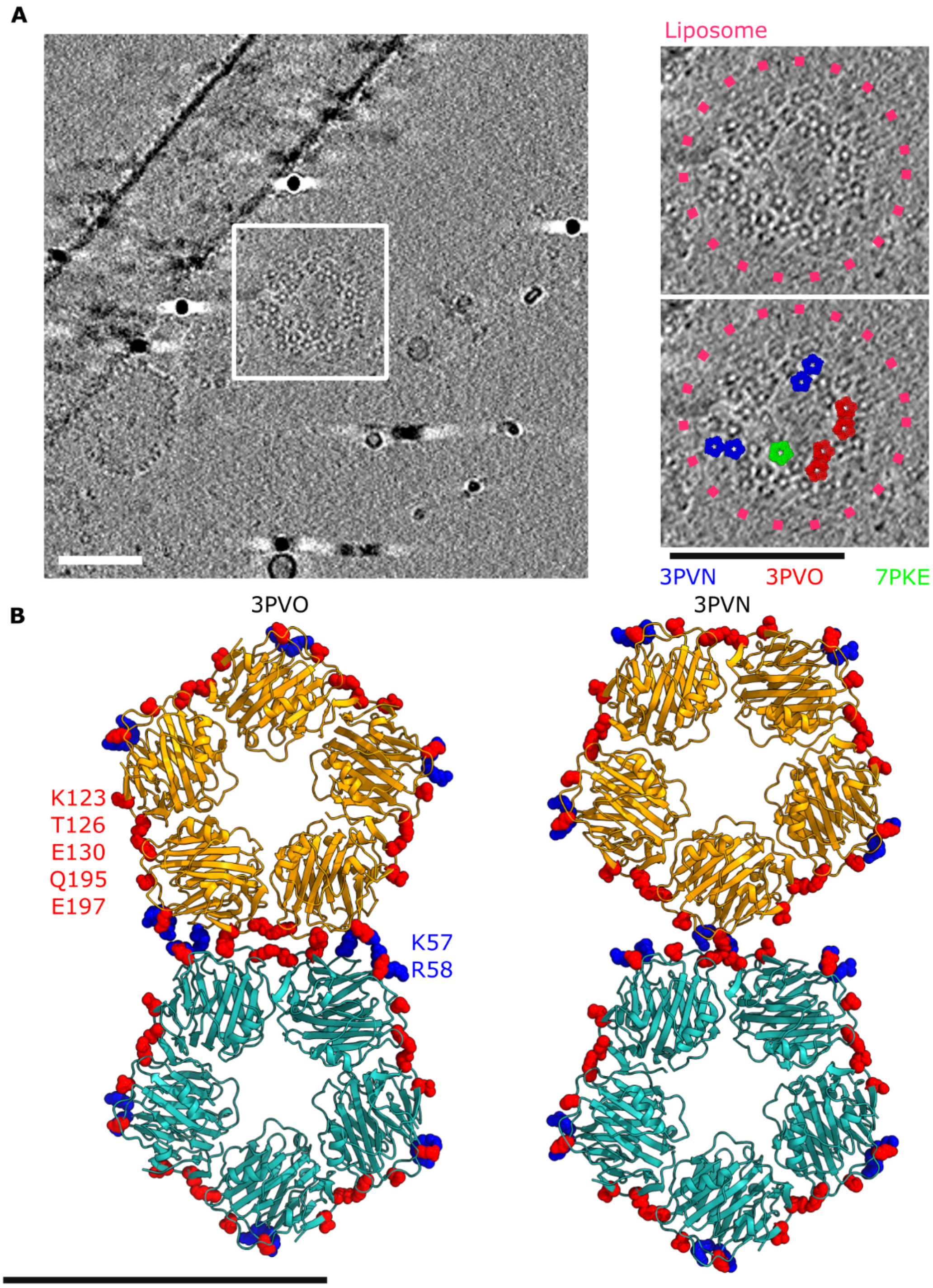


**Figure S11: CRP crystal contacts fit into tomograms of CRP bound to liposomes (DMPC:Cholesterol:DMPG:PE-DBCO, 40:50:5:5 mol%).** (**A**) Slices 14 nm thick through tomograms with 7PKE, 3PVO or 3PVN fitted into density corresponding to CRP. Scale bar represents 100 nm. (**B**) Dimers in the crystal contacts from 3PVO and 3PVN. Residues at lateral regions known to be important in complement activation (blue, K57 & R58) and lateral interactions in the crystal structures (red, K123, T126, E130, Q195 & E197) are highlighted as spheres.

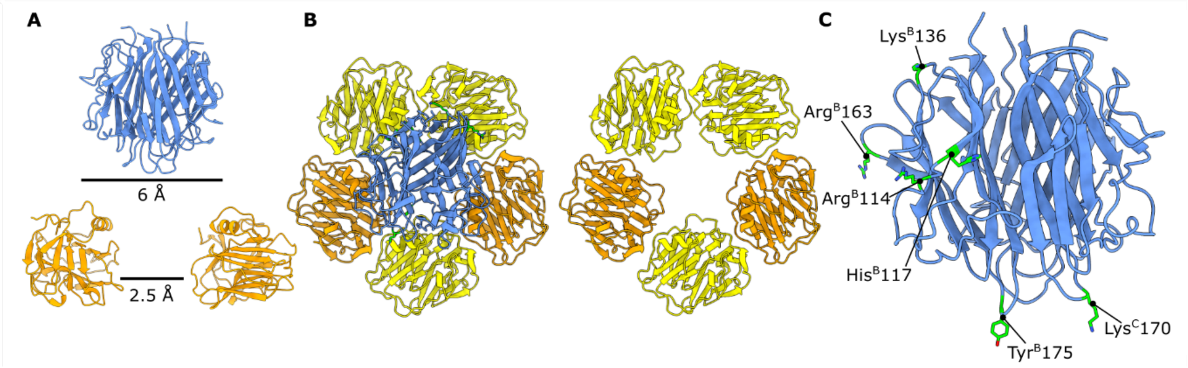


**Figure S12: gC1q interactions with CRP.** (**A**) Dimensions of a single gC1q domain (blue) and the central cavity of the CRP pentamer (orange). A slice through the pentameric toroid of CRP is shown, consisting of two opposing monomers. (**B**) CRP subunits that contact gC1q in the model derived from the subtomogram map, with contacted monomers in yellow and non-contacted in orange. (**C**) Residues on gC1q shown to be important in CRP binding in previous biochemical studies(1) (green).

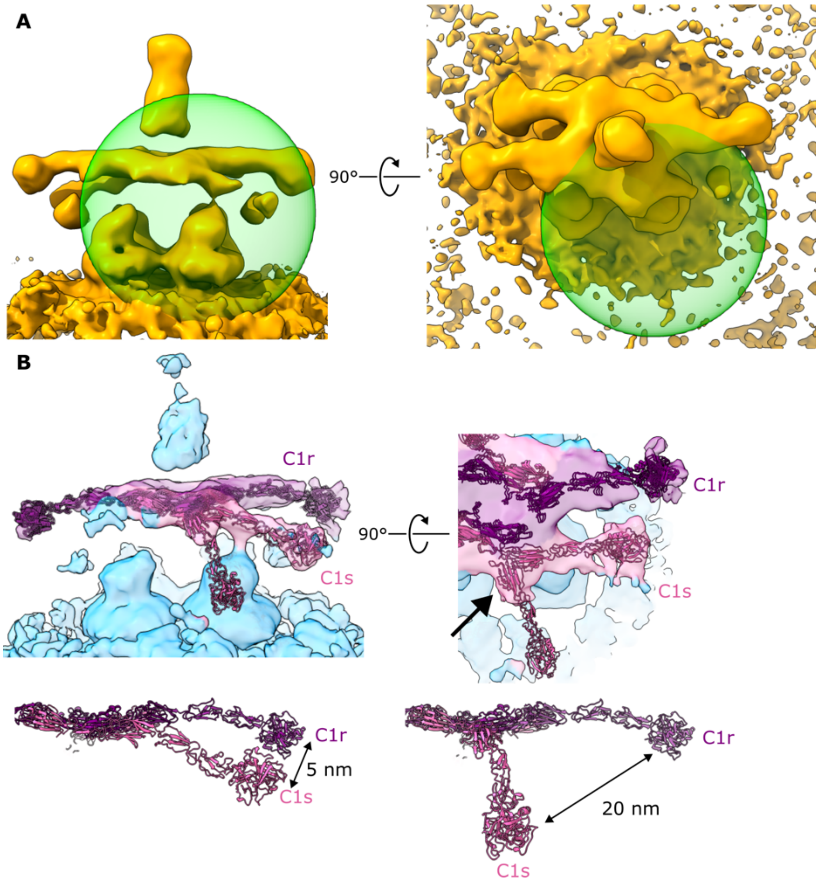


**Figure S13: Masked refinement to resolve C1s heterogeneity.** (**A**) Masking strategy used to resolve the heterogenous region at the C1s domain. The map is low pass filtered to 25 Å. (**B**) Superposition of C1s arms in the raw map, viewed from the top and side.

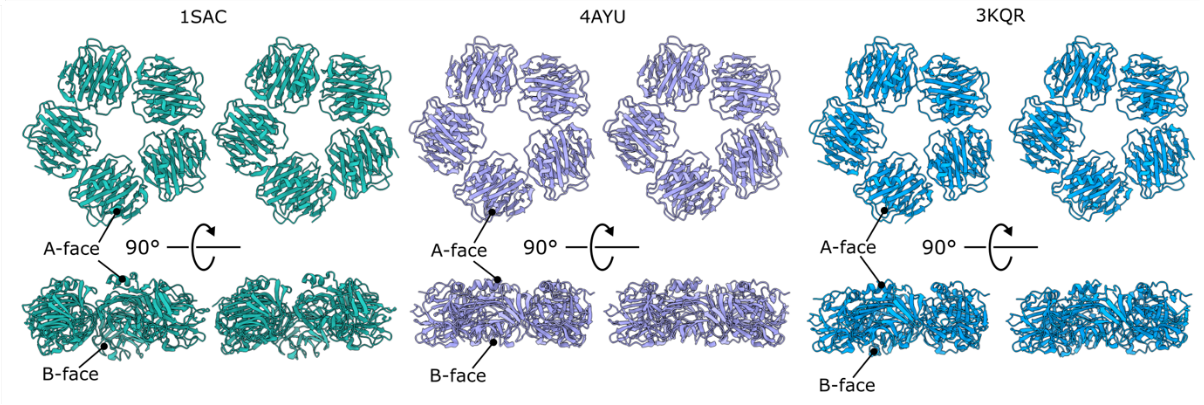


**Figure S14: Crystallographic dimers of laterally interacting pentameric SAP present in the PDB.** Laterally interacting SAP pentamers in a similar arrangement to the CRP dimers from 3PVN and 3PVO were identified in the PDB within the crystal contacts of PDB entries 1SAC, 4AYU and 3KQR.

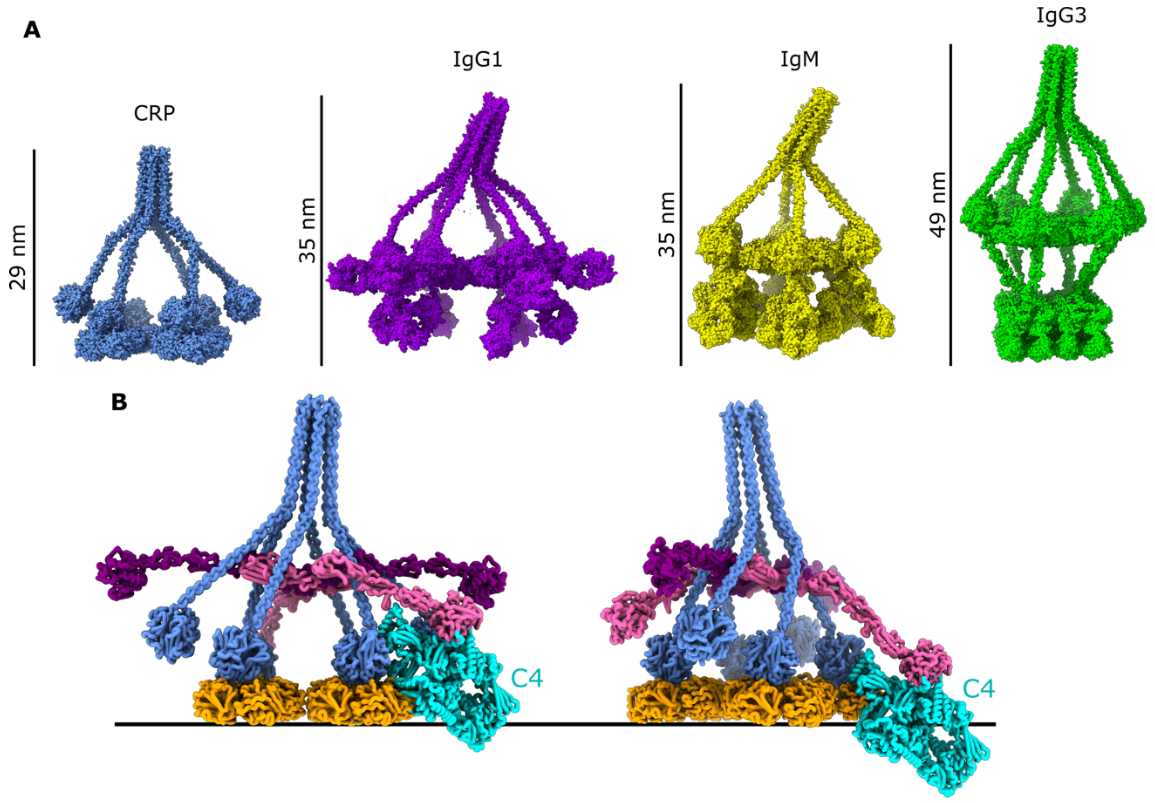


**Figure S15: Comparisons of the heights of C1q-ligand complexes and implications for C4 cleavage.** (**A**) Overall heights of C1 on top of (*l-r*) CRP, IgG1, IgM, and IgG3. (**B**) Aligning the homologous MASP2-C4 complex (PDB code 5JPM(2)) to C1s in either conformation reveals a large steric clash with the lipid membrane (black line).

### Supplementary Methods

#### Liposome binding

First, CRP (1000 nM) was incubated with 150 µl of liposomes for 15 mins at 37°C and 400 RPM. These mixtures were then added to Vivaspin™ spin filters (Sartorius, Germany) with a 300 kDa cut off and made up to 500 µl with TBS, before spinning at 14,000 g at room temperature for 5 minutes. Flow through was discarded and the spin filter was topped up with 500 µl TBS. This was repeated two more times before the material that had not passed through the filter were used for further analysis via SDS-PAGE.

#### Dynamic light scattering

Liposomes (20 µl) were added directly to a DynaPro NanoStar (Wyatt Technology, USA) using a microcuvette (P/N: 162697- 02, 1 µL – DPN, Wyatt technology, USA). The diameter of the particles in solution were measured and technical triplicate was collected with the attenuator set to 50%.

#### C1 proteolysis assay

All steps were done at 4° C until measurements were started with liposomes added at half the total reaction volume. CRP at 2500 nM was added to liposomes (lipid compositions are stated in figure legends)in TBS with 5 mM $CaCl_2$ (reaction buffer). Next, a fluorescent peptide substrate, Boc-Leu-Gly-Arg-Amino Methyl Cumarin (LGR-AMC, PeptaNova GmbH, Germany), dissolved in DMSO (5% DMSO final concentration) was added at 500 µM, before purified C1 complex in reaction buffer (Complement Technology, USA) was added at 40 nM. Subsequently, proteolytic activity of purified C1 complex was measured in real time via cleavage of LGR-AMC on a Clariostar plus plate reader (BMG Labtech, Germany). Measurements were taken every minute for a period of four hours at room temperature, with excitation and emission set at 360 and 460 nm, respectively. The first measurement of each condition was subtracted as background, before normalizing the entire plate to the highest value. Duplicate wells were then averaged and the standard deviation between them was calculated and represented as error bars.

#### *Compound synthesis*

**General methods:** Commercially available solvents and reagents were used as received. Moisture and oxygen sensitive reaction were performed under argon (Ar) atmosphere (balloon). DCM, DMF, ACN and toluene were stored over (flame dried) 4Å molecular sieves. Triethylamine was stored over KOH pellets. TLC analysis was performed using aluminium sheets, pre-coated with silica gel (Merck, TLC Silica gel 60 $F_{254}$). Compounds were visualized with UV absorption (254 nm), by spraying with either 20% $H_2SO_4$ in EtOH, ammonium molybdate/cerium sulphate solution [$(NH_4)_6Mo_7O_{24}·4H_2O$ (25 g/L), $(NH_4)_4Ce(SO_4)_6·2H_2O$ (10 g/L), 10% sulphuric acid in EtOH] or by spraying with aqueous potassium permanganate [$KMnO_4$ (20 g/L), $K_2CO_3$ (10 g/L)]. The purification by column chromatography was carried out with silica gel 60Å (40-63 µm mesh). $^1$H, $^{13}$C and $^{31}$P NMR spectra were recorded on Bruker AV-400 (400-100 MHz) spectrometer with the chemical shifts reported as δ values (ppm). The specta were referenced to TMS (δ=0.00 ppm) in $CDCl_3$ or using the residual solvent peak. The reported coupling constants (*J*) are given in Hz. NMR assignments were made using COSY and HSQC experiments. High resolution mass spectra were recorded by direct injection (2 µL of a 1 µM solution in MeCN) on a Thermo Finnigan LTQ Orbitrap mass spectrometer equipped with an electrospray ion source in positive mode (source voltage 3.5 kV, sheath gas flow 10, capillary temperature 250˚C) with resolution R = 60,000 at m/z 400 (mass range m/z = 150-2,000) and dioctylphthalate (m/z = 391.28428) as a “lock mass”. The HRMS was calibrated prior to measurements with a calibration mixture (Thermo Finnigan).

**2-(2-(2-hydroxyethoxy)ethoxy)ethyl 4-methylbenzenesulfonate (1)**

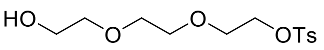


Triethyleneglycol (13.4 mL, 100 mmol) was dissolved in anhydrous DCM (12.5 mL). Triethylamine (2.0 mL, 15.0 mmol) was added. The reaction temperature was lowered to 0°C before adding tosyl chloride (1.91 g, 10.0 mmol) portion wise over 90 min. The reaction mixture was stirred overnight while allowing to reach room temperature. The reaction was diluted with DCM and washed with $H_2O$ (3x). The water layers were back-extracted and the combined organic layers were washed with 10% citric acid (3x), dried over $MgSO_4$, filtrated and concentrated *in vacuo* to obtain **1** (2.98 g, 9.79 mmol, 98%) as a colorless oil. $^1$H NMR (400 MHz, $CDCl_3$) δ 7.81 (d, *J* = 8.3 Hz, 2H, 2x CH Tosyl), 7.35 (d, *J* = 8.3 Hz, 2H, 2x CH Tosyl), 4.17 (t, *J* = 9.6, 4.8 Hz, 2H, H-4), 3.75 – 3.67 (m, 4H, H5-H6), 3.62 (s, 2H, H-1), 3.58 (t, *J* = 9.1, 4.7 Hz, 4H, H3-H4), 2.45 (s, 3H, $CH_3$ Tosyl), 2.24 (t, *J* = 12.1, 6.1 Hz, 1H, OH). $^{13}$C NMR (101 MHz, $CDCl_3$) δ 145.01 ($C_q$ Tosyl), 129.97 (CH Tosyl), 128.11 (CH Tosyl), 72.56, 70.92, 70.44, 69.29, 68.84, 61.89, 21.78 ($CH_3$ Tosyl). HRMS: calculated for $C_{13}H_{20}O_6S$ 322.13188 $[M+NH_4]^+$; found 322.13149.

**2-(2-(2-azidoethoxy)ethoxy)ethan-1-ol (2)**

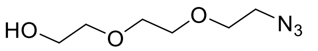


Compound **1** (3.04 g, 10.0 mmol, 1.0 eq) was dissolved in anhydrous DMF (4 mL) and sodium azide (0.65 g, 10.0 mmol, 1.0 eq) was added. The reaction mixture was stirred for 3 h at 90°C. The suspension was filtered and washed with DMF. The residue was concentrated *in vacuo* and co-evaporated with toluene (2x). The crude product was redissolved in DCM and the formed suspension was filtrated and concentrated in vacuo to obtain **2** (1.68 g, 9.59 mmol, 96%) as a yellow oil. $^1$H NMR (400 MHz, $CDCl_3$) δ 3.75 (t, *J* = 8.9, 4.3 Hz, 2H, H-6), 3.72 – 3.66 (m, 6H, H3-H5), 3.65 – 3.60 (m, 2H, H-2), 3.42 (t, *J* = 10.1, 5.3 Hz, 2H, H-1), 2.40 (s, 1H, OH). $^{13}$C NMR (101 MHz, $CDCl_3$) δ 72.59, 70.77, 70.50, 70.18 (C2-C5), 61.88 (C-6), 50.76 (C-1). HRMS: calculated for $C_6H_{13}N_3O_3$ 198.08491 $[M+Na]^+$; found 198.08473.

**Choline-2-cyanoethyl *N,N*-diisopropylphosporamidite tetraphenylborate (3)**

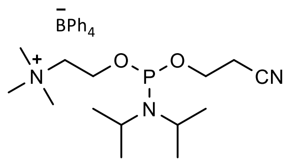


Bis(diisopropylamino)(2-cyanoethoxy)phosphine (0.30 g, 1.0 mmol, 1.0 eq) was dissolved in a mixture of DCM:MeCN (2:1, 3 mL) under Ar atmosphere. Diisopropylammonium tetrazolide (0.17 g, 0.5 mmol, 0.5 eq) and choline tetraphenylborate (0.42 g, 1.0 mmol, 1.0 eq) were added subsequently. The reaction mixture was stirred for 2 h at room temperature. The reaction mixture was diluted with DCM (10 mL) and washed with $NaHCO_3$ (2x 10 mL). Combined aqueous layers back-extracted with DCM. Combined organic layers dried over $MgSO_4$, filtrated and concentrated *in vacuo* obtaining a white solid which was used in the next step without further purification. $^{31}$P NMR (162 MHz, $CDCl_3$) δ 148.1.

**2-(2-(2-Azidoethoxy)ethoxy)ethyl-1-choline phosphate, PC azide (4)**

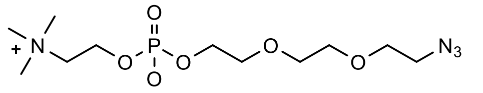


2-(2-(2-Azidoethoxy)ethoxy)ethanol (**2**) was co-evaporated with toluene (2x) and placed under Ar atmosphere. Phosphoramidite **3** (0.52 g, 0.84 mmol, 2.0 eq) dissolved in anhydrous MeCN (10.0 mL) was added. A tetrazole solution in MeCN (0.45 M, 0.93 mL, 0.42 mmol, 1.0 eq) was added and the reaction mixture was stirred for 2 h. Upon completion anhydrous t-BuOOH (5.5 M in nonane, 0.38 mL, 2.10 mmol, 5.0 eq) was added and the reaction was further stirred for 30 min at room temperature ($^{31}$P NMR showed full conversion, $^{31}$P NMR 162 MHz, $CDCl_3$, δ -1.96). DBU (0.31 mL, 2.10 mmol, 5.0 eq) was added and the reaction mixture was stirred for 1 h at room temperature. The reaction mixture was concentrated *in vacuo,* co-evaporated with toluene (2x) and purified with gel-filtration Toyopearl HW40S, MilliQ + 1% AcOH) to obtain **4** (6.0 mg, 17,6 µmol, 4.2%) as a white solid. $^1$H NMR (400 MHz, MeOD) δ 4.33 – 4.26 (m, 2H, H-7), 4.03 – 3.97 (m, 2H, H-6), 3.71 – 3.62 (m, 10H, H2-H5 + H-8), 3.39 (t, *J* = 9.8, 5.1 Hz, 2H, H-1), 3.23 (s, 9H, 3x $CH_3$ choline). $^{13}$C NMR (101 MHz, MeOD) δ 71.88, 71.81, 71.50, 71.48, 71.01, 67.51 (C-8), 67.44 (C-8), 66.15 (C-6), 66.09 (C-6), 60.40 (C-

7), 60.35 (C-7), 54.74 (C-9), 54.70 (C-9), 54.66 (C-9), 51.79 (C-1). $^{31}P$ NMR (162 MHz, MeOD) δ 0.99. HRMS: calculated for $C_{11}H_{25}N_4O_6P$ 341.15845 $[M+H]^+$; found 341.15810.

**NMR spectra PC-azide (4)**

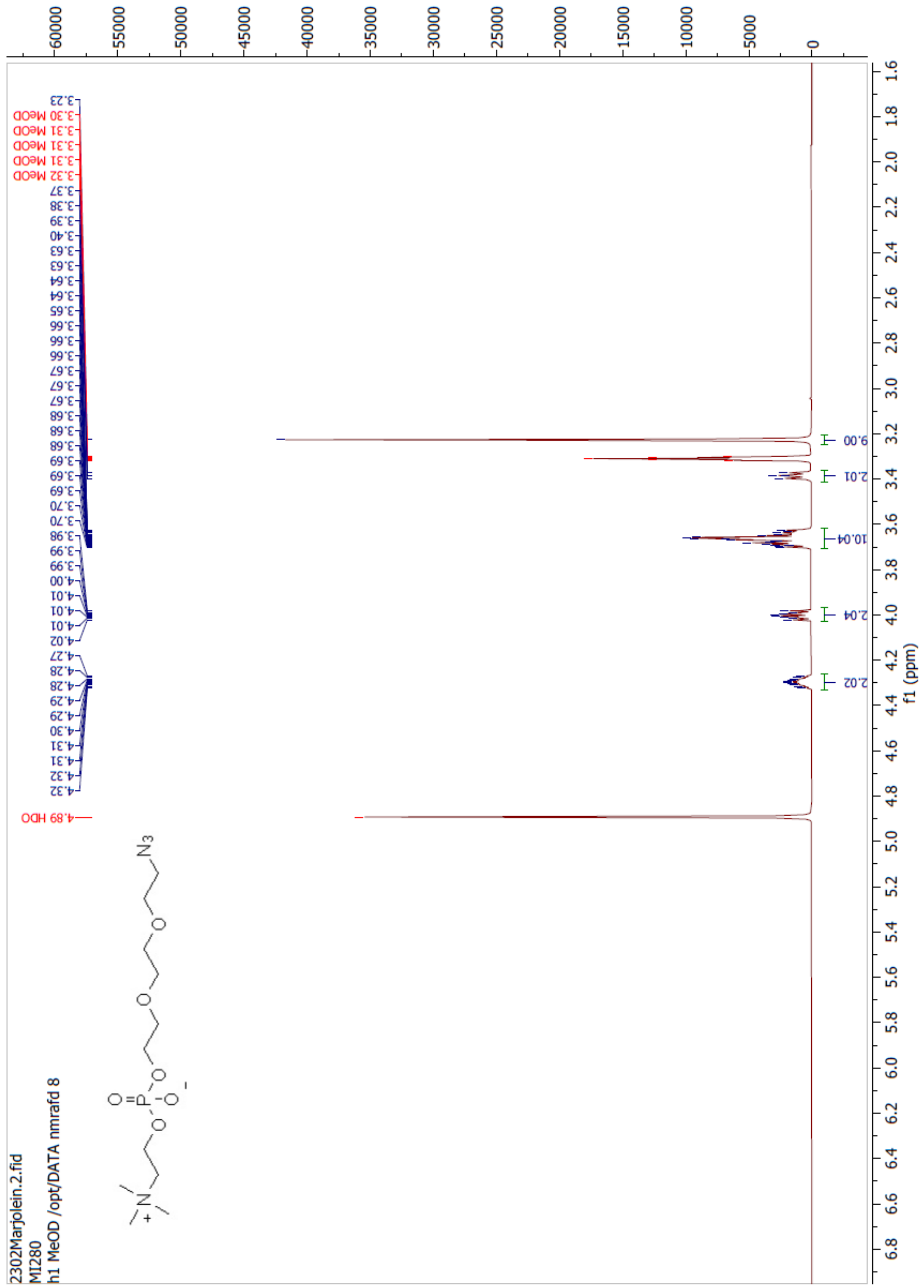

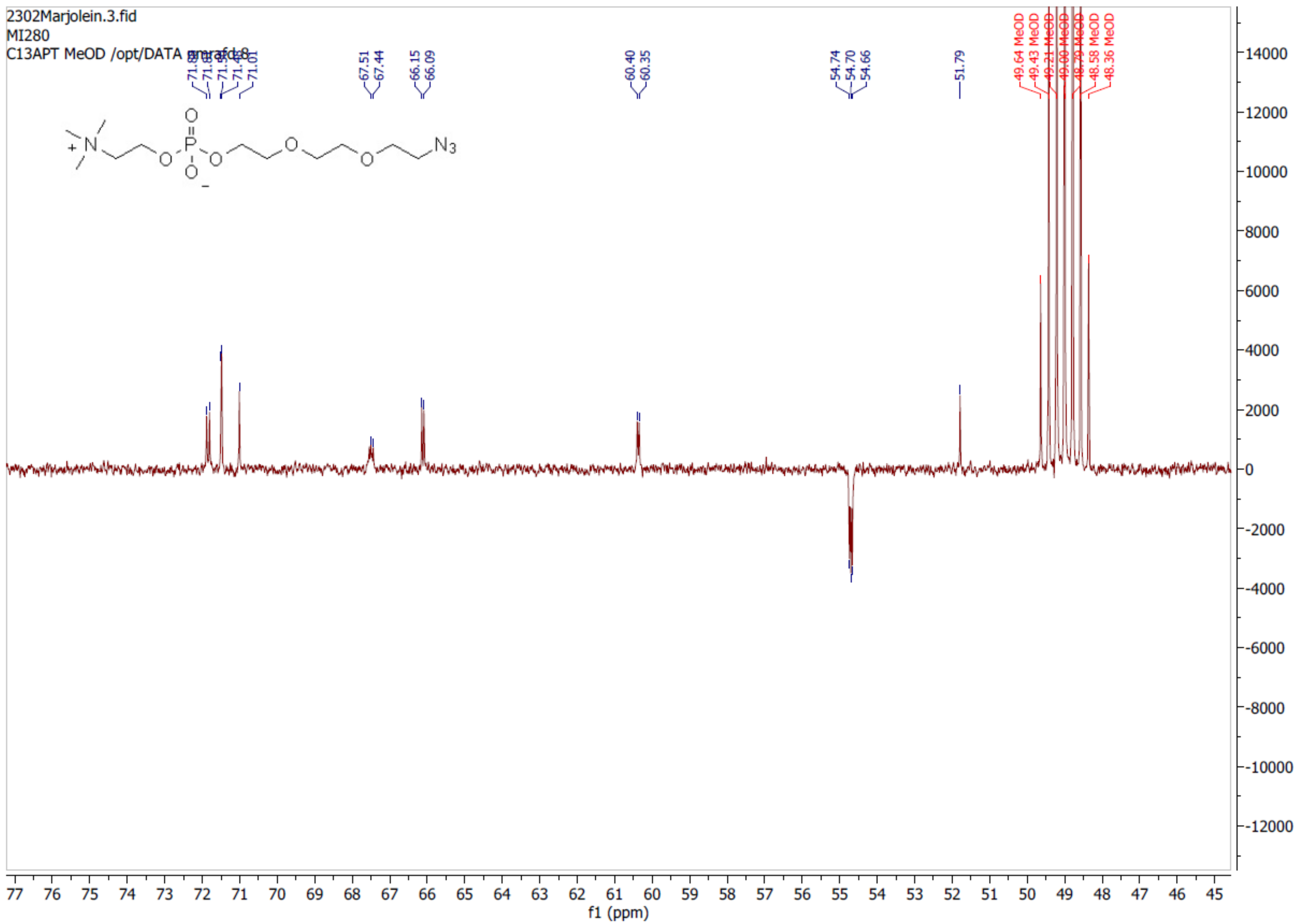
2302Marjolein.3.fid
MI280
C13APT MeOD /opt/DATA
71.01
67.51
67.44
66.15
66.09
60.40
60.35
54.74
54.70
54.66
51.79
49.64 MeOD
49.43 MeOD
49.21 MeOD
49.00 MeOD
48.79 MeOD
48.58 MeOD
48.36 MeOD
N3
f1 (ppm)

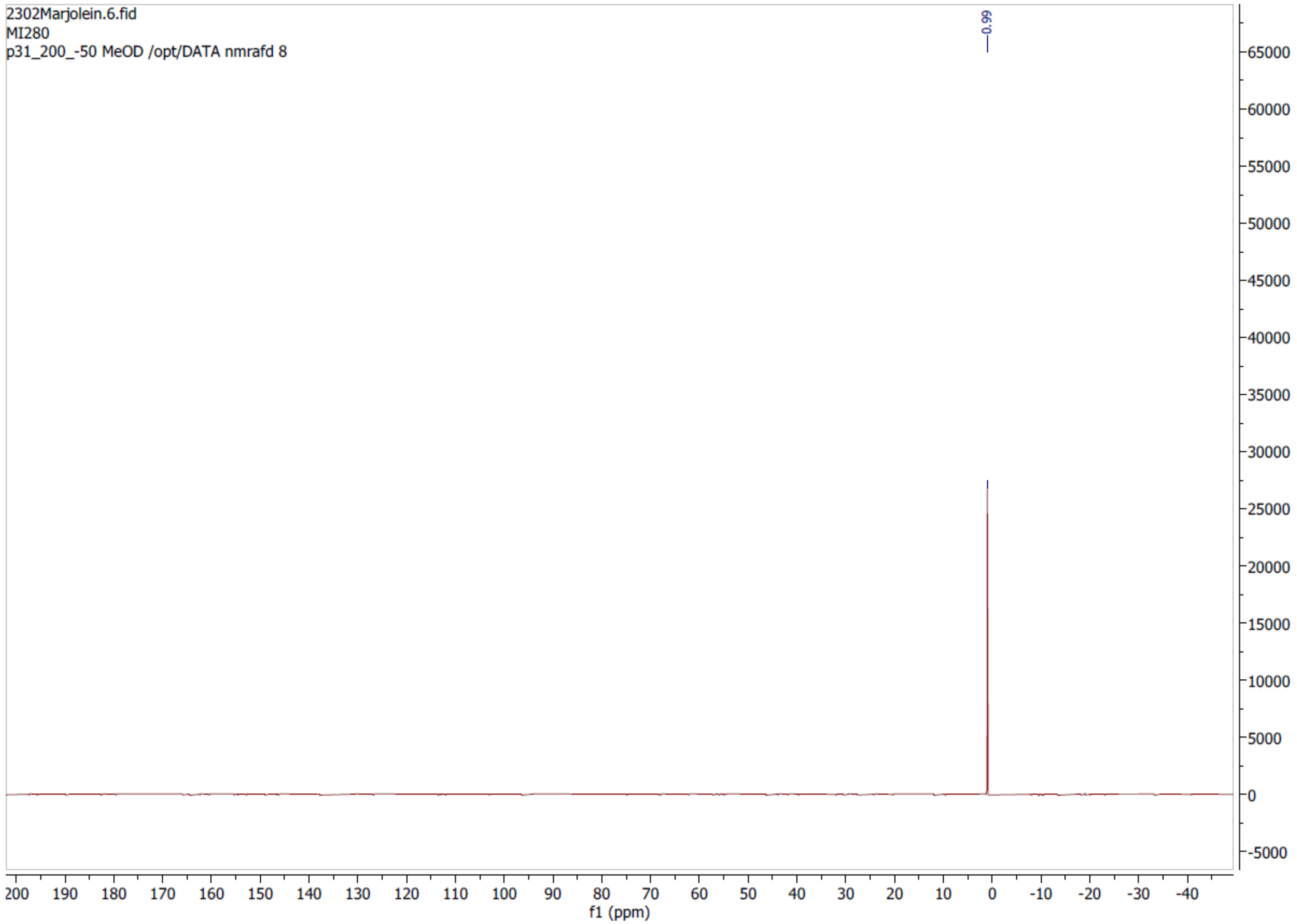
2302Marjolein.6.fid
MI280
p31_200_-50 MeOD /opt/DATA nmrafd 8
0.99
65000
60000
55000
50000
45000
40000
35000
30000
25000
20000
15000
10000
5000
0
-5000
200 190 180 170 160 150 140 130 120 110 100 90 80 70 60 50 40 30 20 10 0 -10 -20 -30 -40
f1 (ppm)
